# Microglia dysfunction caused by the loss of Rhoa disrupts neuronal physiology and leads to neurodegeneration

**DOI:** 10.1101/218107

**Authors:** Renato Socodato, Camila C. Portugal, Teresa Canedo, Artur Rodrigues, Tiago O. Almeida, Joana F. Henriques, Sandra H. Vaz, João Magalhães, Cátia M. Silva, Filipa I. Baptista, Renata L. Alves, Vanessa Coelho-Santos, Ana Paula Silva, Roberto Paes-de-Carvalho, Ana Magalhães, Cord Brakebusch, Ana M. Sebastião, Teresa Summavielle, António F. Ambrósio, João B. Relvas

## Abstract

Nervous tissue homeostasis requires regulation of microglia activity. Using conditional gene targeting in mice, we demonstrate that genetic ablation of the small GTPase Rhoa in adult microglia is sufficient to trigger spontaneous microglia activation producing a neurological phenotype (including synapse and neuron loss, impairment of LTP, formation of ß-amyloid plaques and memory deficits). Mechanistically, loss of Rhoa in microglia triggers Src activation and Src-mediated Tnf production, leading to excitotoxic glutamate secretion. Inhibiting Src in microglia Rhoa-deficient mice attenuates microglia dysregulation and the ensuing neurological phenotype. We also found that the Rhoa/Src signaling pathway was disrupted in microglia of the APP/PS1 mouse model of Alzheimer’s disease and that low doses of Aß oligomers triggered microglia neurotoxic polarization through the disruption of Rhoa-to-Src signaling. Overall, our results indicate that disturbing RhoGTPase signaling in microglia can directly cause neurodegeneration.

## Introduction

Microglia are yolk sack-derived myeloid cells that populate the early-developing nervous system (Ginhoux et al., 2010, Kierdorf et al., 2013, Schulz et al., 2012). Under physiological conditions, microglia continuously extend and retract their cellular processes, monitoring the central nervous system (CNS) parenchyma for tissue damage or infections and checking the functional status of brain synapses (Crotti and Ransohoff, 2016). Although attempting to repair injuries to the parenchyma, exacerbation or prolonged microglial activation may trigger/accelerate neuronal damage in neurogenerative conditions or delay CNS recovery after insults (Gomez-Nicola et al., 2013, Rice et al., 2015, Spangenberg et al., 2016). Such detrimental role for microglia can be attributed to the loss/disruption of microglia immune restrain mechanisms (Deczkowska et al., 2018), which leads to sustained deregulation of their activation and consequent overproduction of proinflammatory cytokines (IL-1β and Tnf), nitric oxide (NO), reactive oxygen species (ROS) and increased release of glutamate (Block et al., 2007).

Proinflammatory microglial activation is associated with profound changes in microglia morphology requiring cytoskeleton reorganization (abd-el-Basset and Fedoroff, 1995, Kaur, 1997). The GTPases of the Rho family, from which Rhoa, Rac1 and Cdc42 are the founding members, are key orchestrators of cytoskeleton dynamics (Bustelo et al., 2007) and are therefore well positioned to govern microglia activation. Typical GTPases function as binary switches by cycling between a GDP-bound inactive, “off” state, and a GTP-bound active, “on” state, in which they can directly interact with downstream effectors to regulate different cell functions (Hodge and Ridley, 2016). RhoGTPases play important roles during CNS development and dysregulation of their expression and/or activity is associated with different neurological disorders (Stankiewicz and Linseman, 2014). Indeed, changes in Rhoa activity or in the activities of its effectors are implicated in several neurodegenerative conditions, including stroke, Alzheimer’s and Parkinson’s Disease and amyotrophic lateral sclerosis (DeGeer and Lamarche-Vane, 2013, Droppelmann et al., 2014). Here, using tamoxifen-inducible conditional gene inactivation in mice, we investigated the bona fide function of Rhoa in microglia. Our data reveal critical roles for Rhoa signaling in regulating microglia immune activity and microglia-dependent synaptic integrity.

## Results

### Conditional ablation of Rhoa in adult microglia

To study the role of Rhoa in adult microglia, we crossed Cx3cr1^CreER-IRES-EYFP^ mice (Parkhurst et al., 2013, Goldmann et al., 2013, Yona et al., 2013) with mice carrying Rhoa conditional alleles (Herzog et al., 2011, Jackson et al., 2011) (**Suppl. Fig. 1A**). In Cx3cr1^CreER^ mice, the CreER-IRES-EYFP transgene is transcriptionally active in Iba1^+^ brain myeloid cells (Goldmann et al., 2016) (**Suppl. Fig. 1B**) and microglia (**Suppl. Fig. 1C**). As expected, following tamoxifen (TAM) administration, Cre-mediated recombination substantially decreased Rhoa expression in Rhoa^fl/fl^:Cx3cr1^CreER+^ microglia (**Fig. 1A and Suppl. Fig. 1D**). Because Rhoa^fl/fl^:Cx3cr1^CreER+^ mice have only one intact copy of Cx3cr1, and decreased Cx3cr1 protein levels could *per se* disrupt microglia homeostasis and be a confounding factor affecting the conclusions of our study, we confirmed that the number of microglia expressing Cxc3r1 as well as microglial number did not vary significantly between Rhoa^fl/fl^ and Cx3cr1^CreER+^(**Suppl. Fig 1E**). We also included other controls to further validate our experimental set up (**Suppl. Fig 1F-H**).

**Figure 1.**
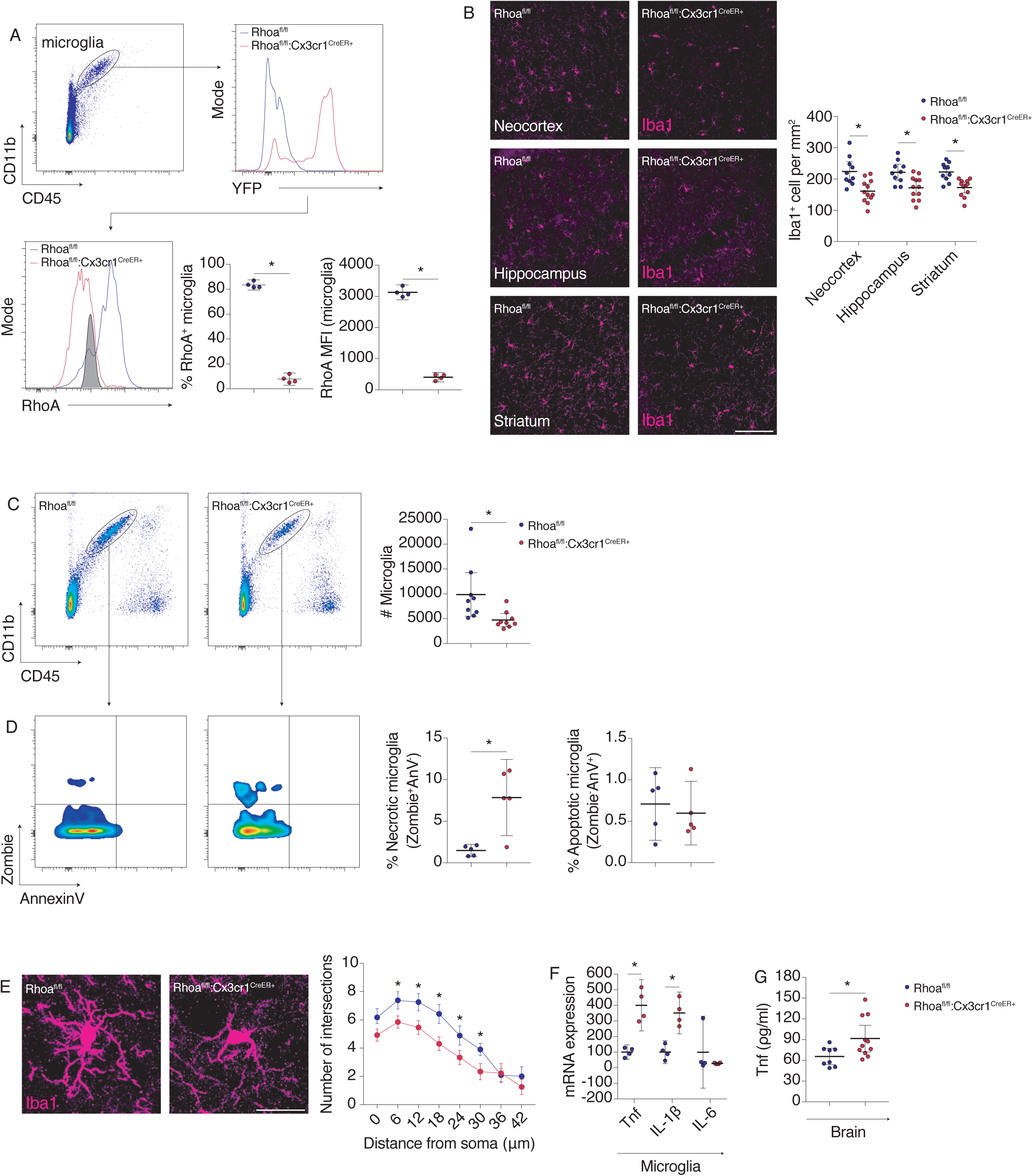
Rhoa is required for microglia survival and its ablation triggers spontaneous microglia activation. **A**, expression of Rhoa in microglia from the brains of Rhoa^fl/fl^ and Rhoa^fl/fl^:Cx3cr1^CreER+^ mice 35-45 days post-TAM (n=4 mice). **B**, confocal images for Iba1 on tissue sections for the indicated brain regions from Rhoa^fl/fl^ and Rhoa^fl/fl^:Cx3cr1^CreER+^ 35-45 days post-TAM (n=10-12 mice). Scale bars: 50 μm. **C**, Flow cytometry analysis of microglial numbers and cell death in Rhoa^fl/fl^ and Rhoa^fl/fl^:Cx3cr1^CreER+^ mice 35-45 days post-TAM (n=9 mice). **D**, Flow cytometry analysis of cell death markers in microglia from Rhoa^fl/fl^ and Rhoa^fl/fl^:Cx3cr1^CreER+^ mice 150 days post-TAM (n=5 mice per condition). **E,** Sholl analysis in microglia from the neocortex of Rhoa^fl/fl^ and Rhoa^fl/fl^:Cx3cr1^CreER+^ 35-45 days post-TAM (n=12-13 cells from 3 mice). Scale bar: 20 μm. **F,** qRT-PCR in microglia sorted from brains of Rhoa^fl/fl^ and Rhoa^fl/fl^:Cx3cr1^CreER+^ mice 35-45 days post-TAM (n=4 mice). **G**, Tnf amounts (ELISA) in Rhoa^fl/fl^ and Rhoa^fl/fl^:Cx3cr1^CreER+^ brains 35-45 days post-TAM (n=8-11 mice). Data are represented as mean with 95% CI. *P<0.05 (Mann-Whitney test in A, C, D, F, G; 2-Way ANOVA in B, E). See also Figures S1-3.

### Rhoa is required for microglial survival and activation

We then evaluated the requirement of Rhoa for microglia homeostasis. We observed a decrease in the number of Iba1^+^ cells in the neocortex, hippocampus and striatum comparing Rhoa^fl/fl^ with Rhoa^fl/fl^:Cx3cr1^CreER+^ littermates (**Fig. 1B**). Using flow cytometry we confirmed a decrease in the numbers of microglia in Rhoa^fl/fl^:Cx3cr1^CreER+^ brains (**Fig. 1C**). Such decrease in microglial numbers persisted in brains of mature adult Rhoa^fl/fl^:Cx3cr1^CreER+^ mice (**Suppl. Fig. 2A**). The decrease on microglial numbers following Rhoa ablation could be caused by increased microglial cell death. Indeed, we found increased necrosis (Zombie^+^Anv^-^) in Rhoa^fl/fl^:Cx3cr1^CreER+^ microglia (**Fig. 1D**). However, microglial numbers in Rhoa^fl/fl^:Cx3cr1^CreER+^ mice 40 days post-TAM (**Fig. 1C**) and 150 days post-TAM (**Suppl. Fig. 2A**) were comparable. This suggested that proliferation could be counterbalancing cell death to stabilize microglial numbers. Accordingly, we found increased microglial proliferation in mature Rhoa^fl/fl^:Cx3cr1^CreER+^ mice compared with Rhoa^fl/fl^ littermates (**Suppl. Fig. 2B**).

Changes in microglial cell morphology are usually concurrent with alterations in microglial function. Morphological analysis revealed that Rhoa mutant microglia were less ramified than Rhoa^fl/fl^ microglia (**Fig. 1E and Suppl. Fig. 2C**). Modifications in actin cytoskeleton organization and dynamics most likely underlie the changes found in microglia morphology. Inhibiting Rhoa activation in microglial cultures by expressing a dominant negative Rhoa mutant (Rhoa^T19N^; **Suppl. Fig. 2D**), resulted in loss of actin stress fibers (**Suppl. Fig. 2E**) and decreased the speed of microglia protrusion extension, which is regulated by actin dynamics (**Suppl. Fig. 2F**).

Next, we evaluated whether production of inflammatory cytokines was increased in Rhoa^fl/fl^:Cx3cr1^CreER+^ microglia. Indeed, Rhoa mutant microglia contained higher amounts of mRNA transcripts coding for the classical proinflammatory cytokines Tnf and IL-1β, but not for IL-6 (**Fig. 1F**). The increased production of Tnf in Rhoa mutant microglia resulted in increased Tnf amounts in Rhoa^fl/fl^:Cx3cr1^CreER+^ brains (**Fig. 1G**).

In addition to microglia, long-lived non-parenchymal macrophages residing on brain interfaces are potentially targeted in Cx3cr1^CreER+^ mice (Goldmann et al., 2016). However, and in contrast to microglia, Rhoa expression in brain macrophages (CD11b^+^CD45^high^ cells; gated as shown in **Suppl. Fig. 2G**) as well as their numbers, viability and reactivity were comparable between Rhoa^fl/fl^:Cx3cr1^CreER+^ mice and aged-matched Rhoa^fl/fl^ littermates (**Suppl. Fig. 2G and H**). Because Cx3cr1 is also expressed on peripheral organs (Jung et al., 2000), some degree of Cre-mediated recombination was expected to occur outside the CNS of Rhoa^fl/fl^:Cx3cr1^CreER+^ mice (Parkhurst et al., 2013). However, blood monocytes and spleen macrophages (**Suppl. Fig. 2I**) showed similar abundance of Rhoa mRNA transcripts between Rhoa^fl/fl^:Cx3cr1^CreER+^ mice and Rhoa^fl/fl^ littermates (**Suppl. Fig. 2J**). The percentage of different CD45^+^ myeloid populations was also comparable between the genotypes (**Suppl. Fig. 2K and L**), indicating that the Rhoa mutant phenotype relates predominantly to microglia and not to macrophages.

To further establish that the loss of Rhoa in microglia would be sufficient to trigger their activation spontaneously in a cell autonomous manner, we depleted Rhoa in different cell culture experimental setups and evaluated microglia activation. We found that (i) loss of Rhoa in primary cortical microglial cultures obtained from Rhoa floxed mice (**Suppl. Fig. 3A**) led to increased expression and secretion of Tnf (**Suppl. Fig. 3B and C**), and that (ii) knocking down (KD) Rhoa in microglial cell lines (**Suppl. Fig. 3D**) also increased Tnf expression (**Suppl. Fig. 3E**) and secretion (**Suppl. Fig. 3F**), (iii) increased NFκB activation (**Suppl. Fig. 3G and H**), and (iv) led to the degradation of the inhibitor of κB (**Suppl. Fig. 3I**) (Shcherbakova et al., 2016).

### Ablation of Rhoa in microglia leads to a neurological phenotype

Increased secretion of Tnf by microglia can be detrimental to neurons because Tnf autocrinally increases the release of glutamate (Takeuchi et al., 2006) thus causing excitotoxic damage to neurites and neurons (Maezawa and Jin, 2010). Using the glutamate release FRET biosensor FLIPE600n^Surface^, which specifically discriminates glutamate release over its uptake or its intracellular mobilization via cell metabolism (Okumoto et al., 2005), we showed that Rhoa KD increased glutamate release from living primary cortical microglia (**Fig. 2A**). This increased release led to an accumulation of glutamate in the extracellular space (**Fig. 2B**). Incubation of primary hippocampal neurons with conditioned media (CM) collected from primary cortical microglia knocked down for Rhoa (Rhoa KD MCM) increased neurite beading (**Fig. 2C**), a characteristic of glutamate excitotoxicity (Maezawa and Jin, 2010). Confirming that microglia-secreted glutamate was driving neurite damage, pharmacological inhibition of the NMDA-type of glutamate receptors on hippocampal neurons abrogated the neurite beading effect of Rhoa KD MCM (**Fig. 2C**).

**Figure 2.**
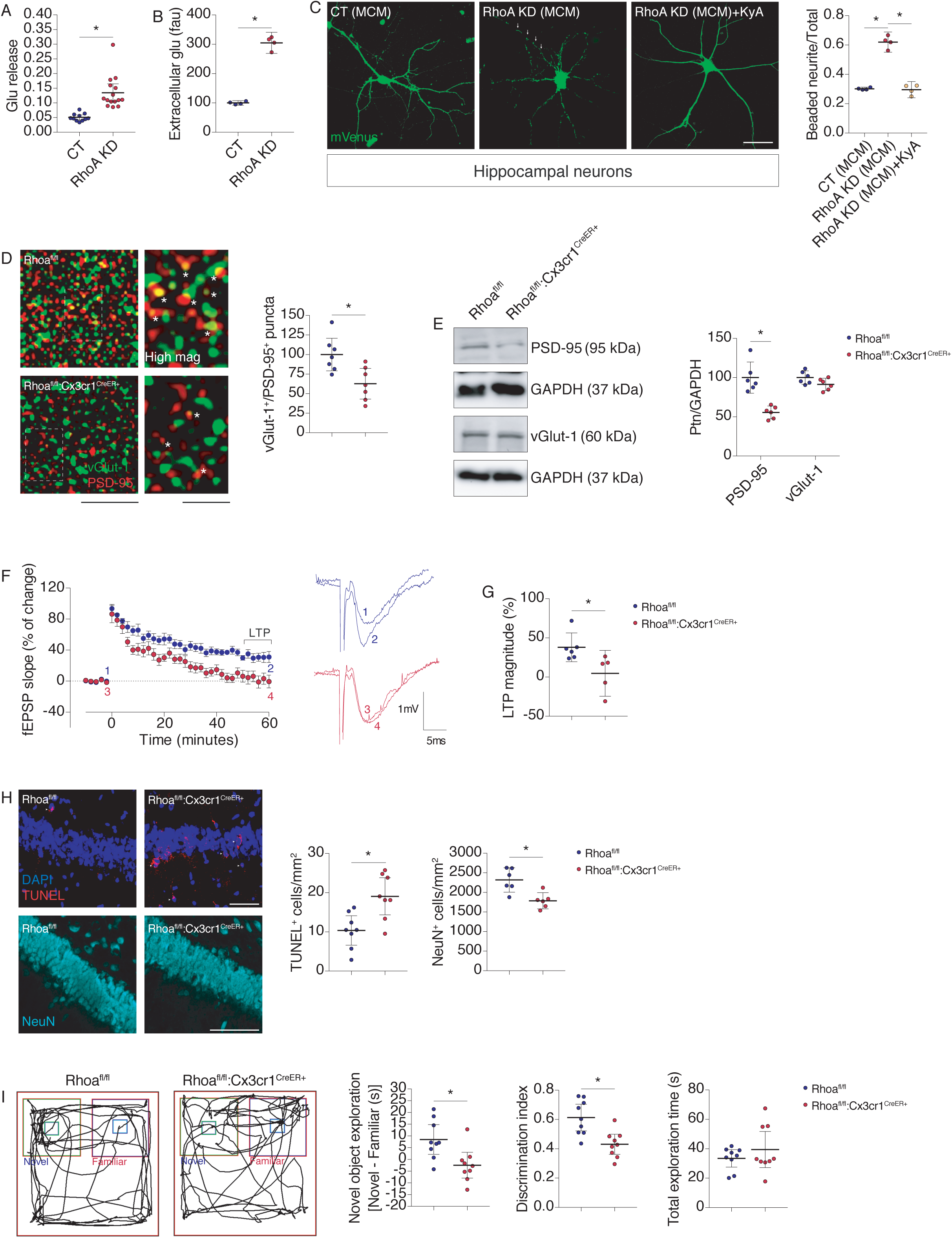
Rhoa deficiency in adult microglia leads to phenotypic features underlying neurological disorders. **A**, primary cortical microglial cultures expressing FLIPE^600nSurface^ were transduced with lentiviruses coding pLKO or Rhoa shRNA (CT; n=11-15 cells from 3 cultures). **B**, Extracellular glutamate in the culture media ofp LKO and Rhoa KD primary cortical microglia cultures (n=4 cultures). fau = fluorescence arbitrary units. **C**, primary hippocampal neuronal cultures expressing mVenus were incubated for 24 h with conditioned media from primary microglial cultures (MCM) obtained from WT mice that were previously infected with pLKO or Rhoa shRNA lentiviruses (n=3-4 cultures). In some conditions neurons were co-incubated with the NMDA receptor antagonist kynurenic acid (KyA; 1 μM). Arrowheads display a beaded neurite. Scale bar: 50 μm. **D**, vGlut-1 (green) and PSD-95 (red) immunolabeling on hippocampus CA1 region of Rhoa^fl/fl^ and Rhoa^fl/fl^:Cx3cr1^CreER+^ mice 35-45 days post-TAM (n=7 mice). Scale bars: 5 μm and 1 μm (high mag). White asterisks denote synaptic puncta. **E,** Western blot for PSD-95 and vGlut-1 on brains lysates from Rhoa^fl/fl^ and Rhoa^fl/fl^:Cx3cr1^CreER+^ mice 35-45 days post-TAM (n=6 mice). GAPDH was the loading control. **F and G**, panel (F) shows the averaged time course changes in field excitatory post-synaptic potential (fEPSP) slope induced by θ-burst stimulation in Rhoa^fl/fl^ (n=11 slices from 6 mice) and Rhoa^fl/fl^:Cx3cr1^CreER+^ (n=9 slices from 5 mice) 35-45 days post-TAM. Ordinates represent normalized fEPSP slopes, from which 0% corresponds to slopes averaged and recorded for 10 min before θ-burst stimulation ((−0.99±0.04 mV/ms (Rhoa^flfl^), −0.93±0.04 mV/ms (Rhoa^fl/fl^:Cx3cr1^CreER+^)), and the abscissa represents the time that average begun. Experiments are the average of six consecutive responses obtained for Rhoa^fl/fl^ or Rhoa^fl/fl^:Cx3cr1^CreER+^ mice before (1 or 3) and 58-60 min after (2 or 4) θ-burst stimulation, and is composed by the stimulus artifact, followed by the pre-synaptic fiber volley and the fEPSP. Panel (G) compares the magnitude of LTP obtained in slices from each genotype; the values in ordinates represent the average of the fEPSP recorded 50-60 min after LTP induction, normalized to pre θ-burst values. **H**, confocal images of hippocampi from Rhoa^fl/fl^ and Rhoa^fl/fl^:Cx3cr1^CreER+^ 35-45 days post-TAM (n=8 mice for TUNEL; n=6 mice for NeuN). Scale bars: 100 μm. **I**, Rhoa^fl/fl^ and Rhoa^fl/fl^:Cx3cr1^CreER+^ were evaluated in the NOR test (after a 4 h delay) 35-45 days post-TAM (n=9 mice). Data are represented as mean with 95% CI. *P<0.05 (Mann-Whitney test in A, B, D-I; Kruskal-Wallis test in C). See also Figures S3 and S4.

We then evaluated the impact of microglia-dependent neurotoxicity in Rhoa^fl/fl^:Cx3cr1^CreER+^ mice. Here we focused our analyses primarily on the hippocampus because this brain region is vulnerable to Tnf-induced glutamate excitotoxicity (Yu et al., 2002). We found less excitatory synaptic puncta in Rhoa^fl/fl^:Cx3cr1^CreER+^ hippocampus (evaluated by the number of vGlut-1^+^/PDS-95^+^ colocalization puncta) compared with Rhoa^fl/fl^ littermates (**Fig. 2D**). Whereas the amounts of vGlut-1 protein were comparable, the amounts of PSD-95 were significantly reduced in Rhoa^fl/fl^:Cx3cr1^CreER+^ hippocampus (**Fig. 2E**).

Because synapse loss can adversely affect neuronal plasticity, we assessed long-term potentiation (LTP) of hippocampal synapses in Rhoa^fl/fl^:Cx3cr1^CreER+^ animals. θ-burst stimulation of hippocampal slices from Rhoa^fl/fl^ mice led to an initial enhancement of field excitatory post-synaptic potentials (fEPSP) slope followed by a decrease and stabilization period, but at the end of recording period, fEPSP slope values remained higher than before θ-burst stimulation (**Fig. 2F blue circles**). In Rhoa^fl/fl^:Cx3cr1^CreER+^ slices, however, the same θ-burst stimulation caused only an initial transient increase in fEPSPs slope values (**Fig. 2F red circles**), which decreased progressively towards pre-θ-burst levels. LTP magnitude was significantly lower on hippocampal slices from Rhoa^fl/fl^:Cx3cr1^CreER+^ mice (**Fig. 2G**), indicating a marked LTP impairment.

Changes in synapse number/function may be associated with neuronal cell death (Palop et al., 2006). Indeed, we found an increase in TUNEL^+^ cells in Rhoa^fl/fl^:Cx3cr1^CreER+^ hippocampus (**Fig. 2H**), indicating neuronal apoptosis. In line with this, Rhoa^fl/fl^:Cx3cr1^CreER+^ hippocampi contained less NeuN positive neurons than hippocampi from Rhoa^fl/fl^ littermates (**Fig. 2H**).

Synapse loss and LTP impairment can result in behavioral deficits. To address this possibility, we compared the performance of Rhoa^fl/fl^:Cx3cr1^CreER+^ and Rhoa^fl/fl^ mice in different behavioral paradigms, including elevated plus-maze (EPM) to evaluate anxiety-like behavior; open field (OF) to test general activity levels and gross motor function; and novel object recognition (NOR) to test recognition memory. Both EPM (**Suppl. Fig. 3J**) and OF (**Suppl. Fig. 3K-P**) tests did not show significant differences between genotypes. In the NOR test, however, Rhoa^fl/fl^:Cx3cr1^CreER+^ mice spent less time exploring the novel object (**Fig. 2I**) and their capacity to discriminate the novel object over the familiar object was reduced, indicating deficits in recognition memory (**Fig. 2I**).

Given that Rhoa, Rhob and Rhoc share high degree of similarity (Wheeler and Ridley, 2004), we asked whether depleting Rhob or Rhoc would also lead to spontaneous microglial activation. Contrasting to Rhoa KD, the KD of Rhob or Rhoc (**validations in Suppl. Fig. 4A**), did not alter microglia morphology (**Suppl. Fig. 4B**), ROS production (**Suppl. Fig. 4C**), glutamate release (**Suppl. Fig. 4D**), NFkB nuclear translocation (**Suppl. Fig. 4E**) or Tnf release (**Suppl. Fig. 4F**). Furthermore, CM from Rhob KD or Rhoc KD microglial cultures did not induce neurite beading on hippocampal neurons (**Suppl. Fig. 4G**). To check whether a compensatory Rhoc upregulation could explain the microglia activation triggered by Rhoa KD, we measured Rhoc activity in living Rhoa KD microglia and detected no overt difference in basal Rhoc activation (**Suppl. Fig. 4H**).

### Rhoa restrains Src tyrosine kinase activity in microglia

We next investigated the signaling pathway downstream of Rhoa controlling microglial activation. Csk, the endogenous repressor of Src family of tyrosine kinases (Nada et al., 1991), plays key roles in inflammation (Thomas et al., 2004). Besides, Src, the prototype and founding SFK member, is a key regulator of microglia activation (Socodato et al., 2015b). Therefore, Csk/Src emerged as good candidates for controlling microglial function and we hypothesized that Rhoa might modulate microglia activation via this pathway.

To test this hypothesis, we investigated whether Rhoa deficiency increased Src activity in microglia. Rhoa^fl/fl^:Cx3cr1^CreER+^ microglia contained higher amounts of active Src compared with control microglia (**Fig. 3A**). Src activity was also increased in Rhoa KD N9 microglia and in Rhoa KD CHME3 microglia (**Fig. 3B and C**). To further demonstrate that Src activity is directly linked to Rhoa function, we used the pSicoR vector (Ventura et al., 2004). In the pSicoR vector system, Cre expression causes recombination and excision of both shRNA and EGFP sequences, turning their expression off (**Fig. 3D**). We found that activation of Src did not change by expressing a scrambled shRNA sequence in lenti-CT microglia or in lenti-Cre microglia (**Fig. 3E**), but significantly increased by expressing shRhoa sequences in lenti-CT microglia (**Fig. 3E**). Confirming the Rhoa-dependent Src modulation, expressing shRhoa in lenti-Cre microglia excised shRhoa sequences and Src activation was retained to control levels (**Fig. 3E**).

**Figure 3.**
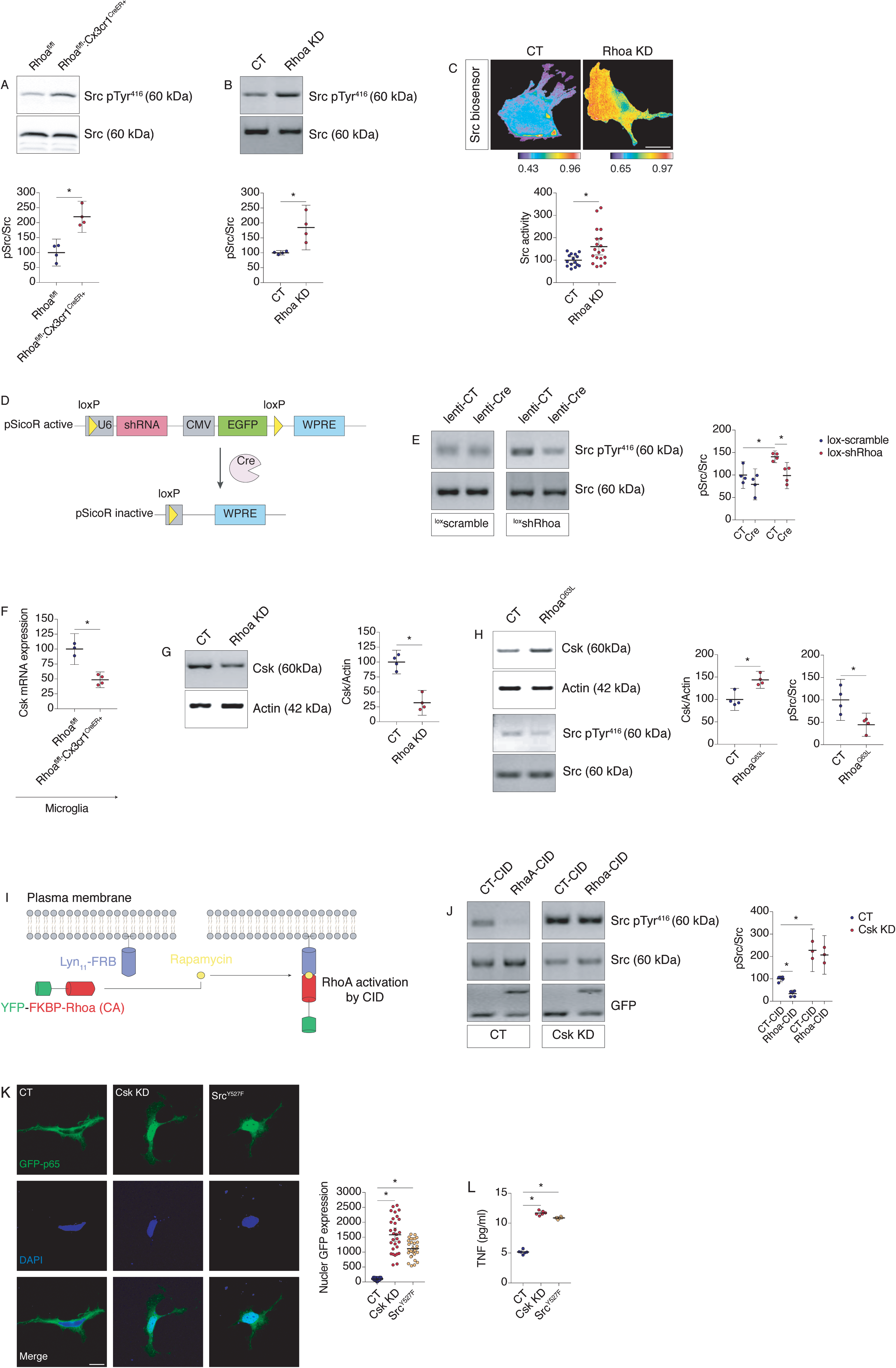
Loss of Rhoa in adult microglia leads to Src hyperactivation through Csk. **A**, Western blot for Src pTyr^416^ on lysates from MACS-separated microglia from Rhoa^fl/fl^ and Rhoa^fl/fl^:Cx3cr1^CreER+^ mice 90 days post-TAM (n=4 mice). Src was the loading control. **B**, Western blot for Src pTyr^416^ on lysates from control (CT) or Rhoa shRNA (Rhoa KD) stable N9 microglial cell sub-clones (n=4 cultures). Src was the loading control. **C**, CHME3 microglial cultures expressing the Src FRET biosensor (KRas Src YPet) were transduced with pLKO or Rhoa shRNA lentiviruses(n=15-19 cells from 3 cultures). Pseudocolor ramps represent min/max CFP/FRET ratios. Scale bar: 5 μm. **D,** schematic representation of pSicoR vector-dependent knockdown of Rhoa (active pSicoR). Rhoa re-expression (inactive pSicoR) is achieved after Cre-dependent recombination of loxP sites. **E**, Western blots for Src pTyr^416^ on lysates from CHME3 microglial cells transduced with lentiviruses carrying pSicoR with a scrambled sequence (lox-scramble-lox) or Rhoa shRNA (lox-shRhoa-lox) (n=4 cultures). In some conditions cells were transfected with an empty vector (lenti-CT) or with a Cre recombinase-expressing vector (lenti-Cre). Src was the loading control. **F,** qRT-PCR on sorted brain microglia from of Cx3cr1^CreER+^ and Rhoa^fl/fl^:Cx3cr1^CreER+^ 35-45 days post-TAM (n=3-4 mice). **G**, Western blot for Csk on lysates from control (CT) or Rhoa shRNA (Rhoa KD) stable N9 microglial cell sub-clones (n=4 cultures). Actin was the loading control. **H**, Western blot for Csk or Src pTyr^416^ on lysates from CHME3 microglial cells transfected with the Rhoa^Q63L^ mutant. Actin and Src were the loading controls (n=4 cultures). **I**, schematic representation of Rhoa activation using chemical-inducible dimerization (CID). **J**, Western blot for Src pTyr^416^ on lysates from pLKO or Csk shRNA stable CHME3 microglial cell sub-clones (n=3-5 cultures). In some conditions cells were co-transfected with YFP-FKBP and Lyn_11_-FRB (CT-CID) or with YFP-FKBP-Rhoa (CA) and Lyn_11_-FRB (Rhoa-CID). All groups were treated with 500 nM rapamycin for 24 h. Src was the loading control. **K,** confocal imaging of p65-GFP-transfected (green) pLKO, Csk KD or Src^Y527F^ stable N9 microglial cell sub-clones (n=29-30 cells from 3 cultures). Nuclei were stained with DAPI (blue). Scale bar: 10 μm. **L,** Tnf amounts (ELISA) were in the culture supernatant of pLKO (CT), Csk KD or Src^Y527F^ stable N9 microglial cell sub-clones (n=6 cultures). Data are represented as mean with 95% CI. *P<0.05 (Mann-Whitney test in A, B, C, F, G, H; Two-Way ANOVA in E, I; One-Way ANOVA in K, L). See also Figure S4.

Csk is the endogenous repressor of Src, and we observed a decrease of Csk mRNA transcripts in Rhoa^fl/fl^:Cx3cr1^CreER+^ microglia (**Fig. 3F**) and of Csk protein amounts in Rhoa KD N9 microglia (**Fig. 3G**). To further link Rhoa activity with regulation of Csk expression, we forced Rhoa activation by expressing the constitutively active Rhoa mutant Rhoa^Q63L^ in microglia. CHME3 microglia expressing Rhoa^Q63L^ had higher amounts of Csk and, consequently, lesser amounts of active Src (**Fig. 3H and Suppl. Fig. 4I and J**), suggesting that Rhoa is an upstream regulator of Csk/Src pathway in microglia.

To strengthen the link between Rhoa activity, Csk expression and Src activity in microglia, we tested whether depleting Csk whilst activating Rhoa would result in Src activation. We forced Rhoa activation using a Rho GTPase chemical inducible dimerization (CID) strategy (**Fig. 3I**) (Inoue et al., 2005). In this experimental set-up the FK506 binding protein (FKBP) and rapamycin binding domain of mTOR (FRB) is fused to the membrane targeting sequence of Lyn (Lyn_11_-FRB) to provide a plasma membrane anchor for the constitutively active form of Rhoa (Rhoa CA), which is expressed as a cytosolic fusion with FKBP and YFP. When microglia expressing both constructs are exposed to rapamycin, YFP-FKBP-Rhoa (CA) rapidly translocates to the plasma membrane and binds to Lyn_11_-FRB triggering fast activation of Rhoa (Inoue et al., 2005) (**Fig. 3I**). Indeed, following exposure to rapamycin, Src activation decreased in pLKO CHME3 microglia co-expressing Lyn_11_-FRB and YFP-FKBP-Rhoa (CA) (Rhoa-CID) relative to microglia co-expressing Lyn_11_-FRB and YFP-FKBP (CT-CID) (**Fig. 3J**). However, in Csk-depleted microglia (Csk KD; **validation in Suppl. Fig 4K**), triggering activation of Rhoa (Rhoa-CID) failed to decrease Src activation (**Fig. 3J**).

Lastly, we examined whether forcing Csk downregulation or Src hyperactivation would be sufficient to activate microglia. Csk KD, which caused a large increase in Src activation (**Suppl. Fig. 4L**), or expressing the constitutively active Src mutant Src^Y527F^ (**Suppl. Fig. 4L; with additional control in Fig. Suppl. Fig. 4M**) led to NFκB activation (**Fig. 3K**) and increased Tnf secretion (**Fig. 3L**) in CHME3 microglia.

### Modulating Rhoa/Src pathway in microglia prevents neurodegeneration

Next, we investigated whether blocking Src would rescue, to some extent, synaptotoxicity and the associated memory deficits in Rhoa^fl/fl^:Cx3cr1^CreER+^ mice. Therefore, following TAM administration, we injected Rhoa^fl/fl^:Cx3cr1^CreER+^ mice weekly with the potent Src blocker AZD 0530 (**Fig. 4A**) and one month later confirmed that this regimen of AZD 0530 was effective in decreasing Src activation in their brain (**Fig. 4B**). A single IP injection of AZD 0530 was sufficient to reduce Src activation in the brains of naïve WT mice (**Suppl. Fig. 5A**) with an apparent IC_50_ of approximately 0.024 mg/kg (**Suppl. Fig. 5B**), indicating that the amounts of AZD 0530 present in the brain after 1 week are sufficient to sustain Src inhibition. Src inhibition by AZD 0530 prevented the decrease of microglial numbers in Rhoa^fl/fl^:Cx3cr1^CreER+^ mice (**Fig. 4C and D; Suppl. Fig. 5C and D**) and the increase of mRNA transcript of Tnf in Rhoa^fl/fl^:Cx3cr1^CreER+^ brains (**Fig. 4E**).

**Figure 4.**
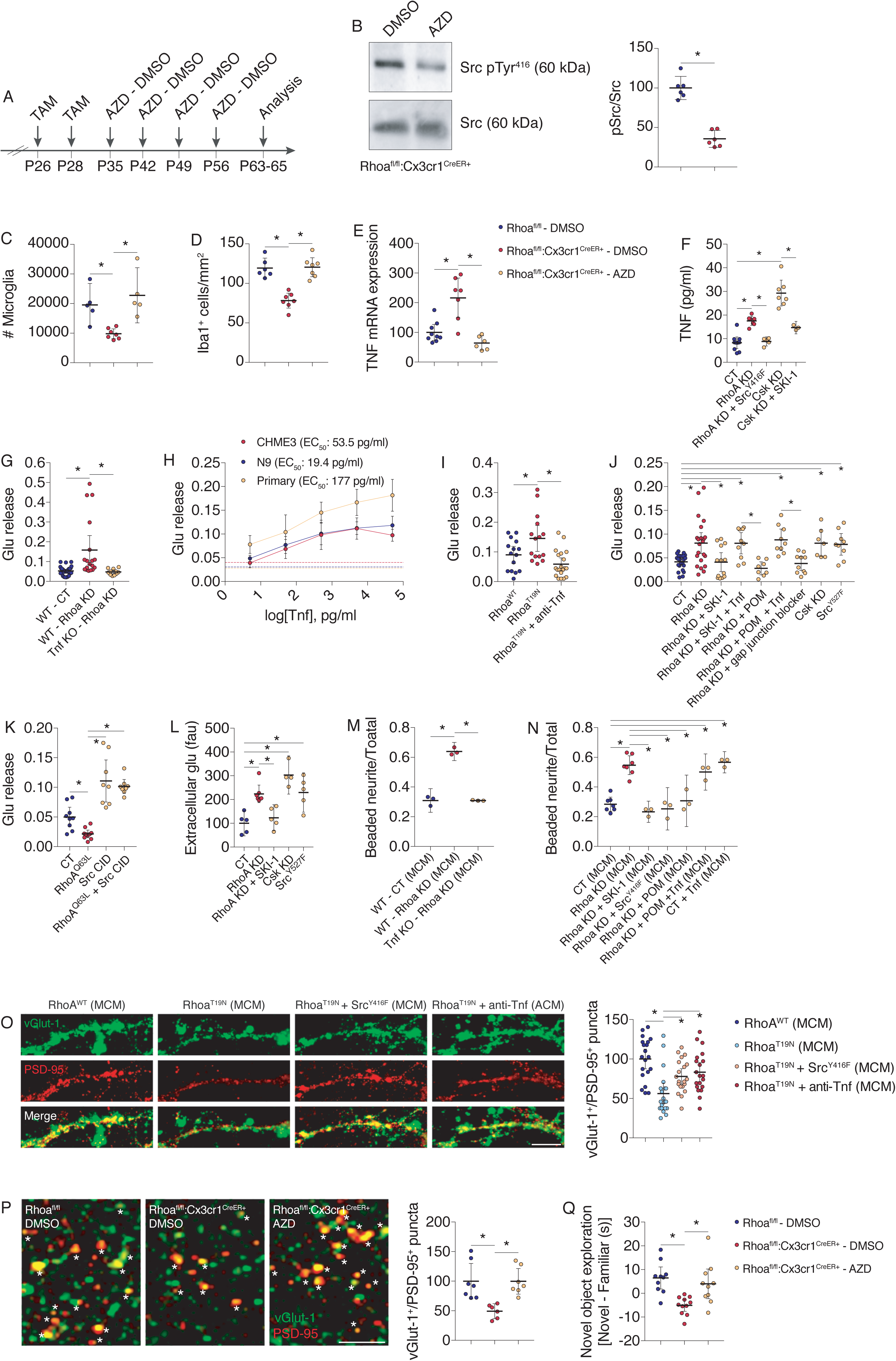
Src inhibition compensates Rhoa deficiency in microglia preventing synapse loss and memory deficits. **A**, regimen of 10 mg/kg AZD 0530 or DMSO injections to Rhoa^fl/fl^:Cx3cr1^CreER+^ mice after TAM administration. **B**, Western blot for Src pTyr^416^ on lysates from the brains of Rhoa^fl/fl^:Cx3cr1^CreER+^ mice injected with 10 mg/kg AZD 0530 or DMSO (n=6 mice). Src was used as the loading control. **C**, FACS analysis of microglia in Rhoa^fl/fl^ and Rhoa^fl/fl^:Cx3cr1^CreER+^ mice injected with 10 mg/kg AZD 0530 or DMSO (n=5-7 mice). **D**, Iba1 immunolabeling on tissue sections from the hippocampal dentate gyrus of Rhoa^fl/fl^ and Rhoa^fl/fl^:Cx3cr1^CreER+^ mice injected with 10 mg/kg AZD 0530 or DMSO (n=6-7 mice). **E**, qRT-PCR from the neocortex of Rhoa^fl/fl^ and Rhoa^fl/fl^:Cx3cr1^CreER+^ mice injected with 10 mg/kg AZD 0530 or DMSO (n=6-9 mice). **F,** Tnf amounts (ELISA) in the culture supernatant of pLKO (CT), Rhoa KD or Rhoa KD:Src^Y416F^, Csk KD or Csk KD treated with SKI-1 (500 nM; 24 h) stable N9 microglial cell sub-clones (n=3-10 cultures). **G,** primary microglia from WT or Tnf KO mice were infected with pLKO (CT), Rhoa KD lentiviruses and then transfected with the glutamate release FRET biosensor FLIPE (n=11-32 cells from 3 cultures). **H**, CHME3 microglia (n=6 cells from 2 independent cultures), N9 microglia (n=6 cells from 2 cultures) or primary microglia (n=4 cells from 2 cultures) expressing the glutamate release FRET biosensor FLIPE were treated with different concentrations of recombinant Tnf. Traced lines represent baseline glutamate release. **I**, CHME3 microglial cells were co-transfected with the Rhoa^WT^ or Rhoa^T19N^ and the glutamate release FRET biosensor FLIPE (n=15-20 cells from 2 cultures). Cultures were then incubated with an FcR blocking solution for 10 min and in some conditions Rhoa^T19N^ cells were treated with Adalimumab (anti-Tnf; 5 μg/ml; 48 h). **J**, pLKO (CT), Rhoa KD, Csk KD or Src^Y527F^ CHME3 microglial cell sub-clones were transfected with the glutamate release FRET biosensor FLIPE (n=7-23 cells from 3 cultures). In some conditions Rhoa KD cells were treated with the Src inhibitor SKI-1 (500 nM; 24 h), the inhibitor of Tnf production pomalidomide (POM; 500 nM; 24 h), recombinant Tnf (30 pg/ml; 24 h) or the gap junction blocker 18-alpha-glycyrrhetinic acid (10 μM; 24 h). **K**, CHME3 microglial cells expressing the glutamate release FRET biosensor FLIPE were co-transfected with pLKO (CT) or Rhoa^Q63L^ together with RapR-Src and FRB-mCherry (n=8-10 cells from 3 cultures) and treated with 500 nM rapamycin for 90 min. **L**, glutamate amounts in the culture media from pLKO (CT), Rhoa KD, Csk KD or Src^Y527F^ N9 microglial cell sub-clones (n=4-7 cultures). In some conditions Rhoa KD cells were treated with the Src inhibitor SKI-1 (500 nM; 24 h). fau = fluorescence arbitrary units. **M**, primary hippocampal neuronal cultures expressing mVenus were incubated for 24 h with MCM obtained from primary microglia cultures from WT mice (infected with CT or Rhoa KD lentiviruses) or Tnf KO mice (infected with Rhoa KD lentiviruses) (n=3 cultures). **N**, primary hippocampal neuronal cultures expressing mVenus were incubated for 24 h with MCM obtained from pLKO (CT), Rhoa KD, and RhoA KD + Src^Y416F^ N9 microglial cell sub-clones (n=3-7 cultures). In some conditions Rhoa KD microglia were treated with SKI-1 (500 nM; 24 h) or pomalidomide (POM; 500 nM; 24 h). Recombinant Tnf (30 pg/ml; 24 h) was used in CT microglial clones. **O**, vGlut-1 (green) and PSD-95 (red) immunolabelling in primary hippocampal neurons incubated for 24 h with conditioned media from CHME3 microglial cultures (MCM) overexpressing Rhoa^WT^, Rhoa^T19N^ or Src^Y416F^ (n=20 neurites from 2 cultures). Microglial cultures were pre-incubated with an FcR receptor blocking solution for 20 min and in some treated with Adalimumab (anti-Tnf; 5 μg/ml; 48 h). Scale bar: 5 μm. **P**, vGlut-1 (green) and PSD-95 (red) immunolabeling on tissue sections from the hippocampal CA1 region of Rhoa^fl/fl^ and Rhoa^fl/fl^:Cx3cr1^CreER+^ mice injected with 10 mg/kg AZD 0530 or DMSO (n=6-7 mice). Scale bar: 5 μm. Asterisks denote synaptic puncta (yellow). **Q**, DMSO and AZD 0530-injected Rhoa^fl/fl^ or Rhoa^fl/fl l^:Cx3cr1^CreER+^ mice were evaluated in the NOR test (n=10-11 mice). Data are represented as mean with 95% CI. *P<0.05 (Mann-Whitney test in B; Kruskal-Wallis test in C-E, P; One-way ANOVAin F, G, I-O, Q). See also Figure S5.

Expressing the dominant negative Src mutant Src^Y416F^ in Rhoa KD microglia prevented Tnf release compared with Rhoa KD microglia (**Fig. 4F**). Src^Y416F^ expression also caused reduced activation of NFκB in Rhoa KD microglia (**Suppl. Fig. 5E**). Pharmacological inhibition of Src, using the Src blocker SKI-1, in Csk KD microglia also prevented Tnf release (**Fig. 4F**) and NFκB nuclear translocation (**Suppl. Fig. 5E**).

Because blocking Src activity in microglia lacking Rhoa abrogated Tnf production we postulated that it would also inhibit glutamate release, thereby preventing microglia from becoming neurotoxic. Aablating microglial Tnf (using Tnf KO) abrogated the release of glutamate triggered by depleting Rhoa (**Fig. 4G**) and Tnf dose-dependently increased the release of glutamate from CHME3 microglia, N9 microglia and primary microglia (**Fig. 4H**). We further confirmed that Tnf released from microglia was inducing the release of glutamate using Adalimumab (a monoclonal anti-Tnf blocking antibody), which blocked the release of glutamate triggered by overexpressing Rhoa^T19N^ in microglia (**Fig. 4I**). Whereas the Src inhibitor SKI-1 attenuated the release of glutamate in Rhoa KD microglia (**Fig. 4J**), exogenous application of Tnf rescued the release of glutamate after inhibiting Src in Rhoa KD microglia (**Fig. 4J**). Preventing Tnf production in Rhoa KD microglia, using the antineoplastic Tnf inhibitor pomalidomide (POM), also inhibited the increase of glutamate release (**Fig. 4J**), which could again be rescued by exogenous application of Tnf (**Fig. 4J**).

One main route for glutamate release from active microglia is its extrusion through gap junction hemichannels (Maezawa and Jin, 2010, Socodato et al., 2018). Blocking gap junctions using 18-alpha-glycyrrhetinic acid in Rhoa KD microglia prevented glutamate release (**Fig. 4J**). In addition, hyperactivating Src (by Csk KD or by expressing Src^Y527F^) recapitulated the effect of Rhoa KD in microglia and increased glutamate release (**Fig. 4J**).

Increasing basal Rhoa activation in microglia by expressing Rhoa^Q63L^ was sufficient to decrease glutamate release (**Fig. 4K**). Therefore, we hypothesized that if Src acted downstream of Rhoa to promote glutamate release, then forcing Src activation, which increases Tnf production, in Rhoa^Q63L^-expressing microglia should result in glutamate release. To test this hypothesis, we used a Src-CID strategy to exert precise temporal control over Src activation (Karginov et al., 2010). Exposing microglial cultures co-expressing RapR-Src and FRB-mCherry (Src-CID) to rapamycin led to a significant increase in glutamate release (**Fig. 4K**) and resulted in the release of glutamate in cells expressing Rhoa^Q63L^ (**Fig. 4K**). Rhoa KD or direct Src hyperactivation led to an accumulation of glutamate in the extracellular medium (**Fig. 4L**). Inhibiting Src with SKI-1 in Rhoa KD microglia also abrogated extracellular accumulation of glutamate (**Fig. 4L**).

We assessed whether inhibiting Src or Tnf after depleting Rhoa would attenuate neurite damage in hippocampal neurons. CM from Tnf-deficient microglia prevented neurite beading caused by Rhoa KD (**Fig. 4M and Suppl. Fig. 5F**). Inhibiting Src in microglia (using SKI-1 or the dominant negative Src mutant Src^Y416F^) prevented neurite beading in cultured hippocampal neurons caused by the CM of Rhoa KD microglia (**Fig. 4N and Suppl. Fig. 5G**). CM from Rhoa KD microglia in which Tnf was inhibited with POM also prevented neurite beading and adding recombinant Tnf to Rhoa KD microglia in the presence of POM induced neurite beading (**Fig. 4N and Suppl. Fig. 5G**). The CM of control microglia treated with Tnf also led to extensive neurite beading (**Fig. 4N and Suppl. Fig. 5G**). Paralleling the effect in neurite damage, the CM from microglia overexpressing Rhoa^T19N^ produced a significant loss of synaptic puncta in primary hippocampal neurons (**Fig. 4O**) and decreasing microglial Src activity or blocking microglial Tnf significantly attenuated this effect (**Fig. 4O**). Accordingly, Rhoa^fl/fl^:Cx3cr1^CreER+^ mice injected with AZD 0530 not only contained more hippocampal excitatory synapses (**Fig. 4P**), but also had increased amounts of the structural synaptic markers synapsin-1, synaptophysin and PSD-95 compared with Rhoa^fl/fl^:Cx3cr1^CreER+^ mice injected with vehicle (**Suppl. Fig. 5H**).

Next, we evaluated, using the NOR test, whether AZD 0530 would also attenuate memory deficits after Rhoa ablation in microglia. Rhoa^fl/fl^:Cx3cr1^CreER+^ mice injected with AZD 0530 spent more time exploring the novel object (**Fig. 4Q**) and discriminated the novel object better than Rhoa^fl/fl^:Cx3cr1^CreER+^ mice injected with vehicle (**Suppl. Fig. 5I**). We observed no differences in total object exploration time between Rhoa^fl/fl^:Cx3cr1^CreER+^ mice injected with AZD 0530 and controls (**Suppl. Fig. 5J**).

We did not find significant differences between DMSO-injected Cx3cr1^CreER+^, AZD-injected Cx3cr1^CreER+^ or Rhoa^fl/fl^ mice in microglial numbers (**Suppl. Fig. 5K**), excitatory synapse number (**Suppl. Fig. 5L**), recognition memory (**Suppl. Figs. 5M and N**), anxiety-related behavior (**Suppl. Figs. 5O-S**) and locomotor activity (**Suppl. Fig. 5T**), suggesting that AZD administration did not affect these features at the steady state.

We also injected Rhoa^fl/fl^:Cx3cr1^CreER+^ and control Cx3cr1^CreER+^ mice with POM (50 mg/Kg IP 3 times per week over 4 weeks) and found that this dosage of POM decreased the expression of Tnf in the brains of Cx3cr1^CreER+^ and Rhoa^fl/fl^:Cx3cr1^CreER+^ mice (**Suppl. Fig. 5U**). Besides decreasing brain Tnf expression, POM administration also significantly prevented the decrease of vGlut-1^+^/PSD95^+^ synapses in Rhoa^fl/fl^:Cx3cr1^CreER+^ mice (**Suppl. Fig. 5V**), suggesting that the increased Tnf expression was, at least in part, involved in the synapse loss elicited by the ablation of Rhoa in microglia.

### Rhoa ablation in microglia produces an amyloid-like pathology

Together with synapse loss, LTP impairment and memory deficits (**Fig. 2**), deposition of amyloid beta (Aβ) underlies multiple CNS neurodegenerative states, including Alzheimer’s Disease (AD). Ablation of Rhoa in microglia resulted in increased amounts of Aβ (**Fig. 5A**) and its amyloidogenic precursor β-CTF (a carboxyl-terminal fragment generated by the cleavage of APP by β-secretases; **Fig. 5B**) in Rhoa^fl/fl^:Cx3cr1^CreER+^ brains. Total APP levels were comparable between genotypes (**Fig. 5B**). Corroborating the increased amyloidogenic processing of APP, we detected increased amounts of Aβ_1-40_ peptide and of Aβ_1-42_ peptide in Rhoa^fl/fl^:Cx3cr1^CreER+^ brains (**Fig. 5C**). We also found that the excess of A β formed fibrillar plaque-like structures (labeled by methoxy-X04) in the neocortex of Rhoa^fl/fl^:Cx3cr1^CreER+^ mice but not in Rhoa^fl/fl^ neocortices (**Fig. 5D**).

**Figure 5.**
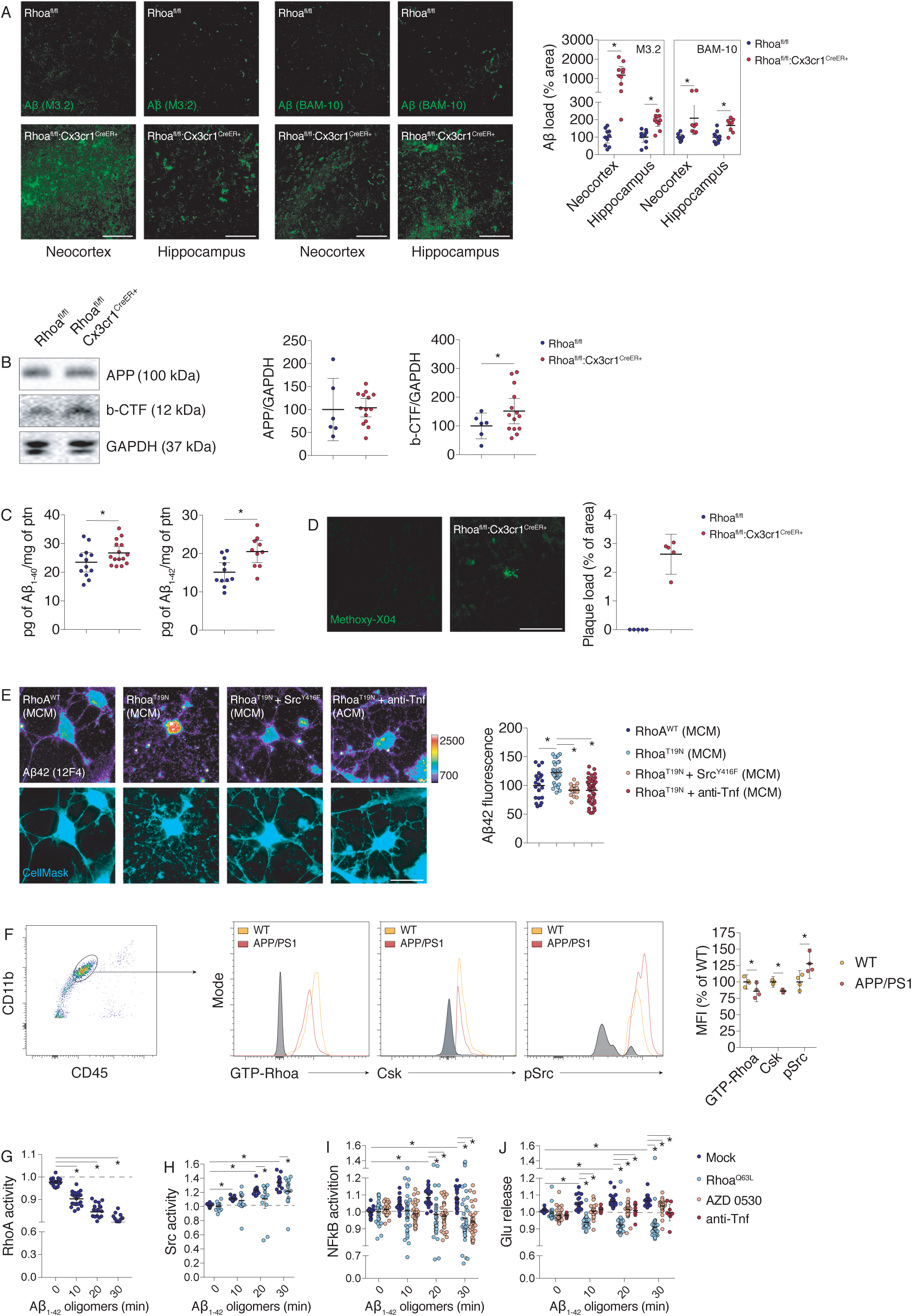
Loss of Rhoa in adult microglia produces an amyloid-like pathology. **A,** confocal images of Aβ deposits in Rhoa^fl/fl^ and Rhoa^fl/fl^:Cx3cr1^CreER+^ mice 50 days post-TAM (n=7 mice for BAM-10 and 10 mice for M3.2). Scale bars: 100 μm. **B**, Western blot using M3.2 antibody (detecting APP and β-CTF) on lysates from the brains of Rhoa^fl/fl^ and Rhoa^fl/fl^:Cx3cr1^CreER+^ 50 days post-TAM (n=6-14 mice). GAPDH was the loading control. **C**, ELISA for Aβ_1-40_ and Aβ_1-42_ on neocortical extracts of RhoA^fl/fl^ and RhoA^fl/fl^:Cx3cr1^CreER+^ 50 days post-TAM (n=12-15 mice for Aβ_1-40_ and n=10-11 mice for Aβ_1-42_). **D**, confocal images of methoxy-X04^+^ plaque-like deposits on RhoA^fl/fl^ and RhoA^fl/fl^:Cx3cr1^CreER+^ neocortices 50 days post-TAM (n=5 mice). **E**, Intracellular endogenous Aβ42 (immunolabelled with anti-Aβ42 clone 12F4) and CellMask dye (cyan) in primary hippocampal neurons incubated for 24 h with conditioned media from CHME3 microglial cultures (MCM) overexpressing Rhoa^WT^, Rhoa^T19N^ or Src^Y416F^ (n=15-40 cells from 2 cultures). Microglial cultures were pre-incubated with an FcR receptor blocking solution for 20 min and in some conditions also treated with Adalimumab (anti-Tnf; 5 μg/ml; 48 h). Scale bar: 20 μm. **F**, flow cytometry showing expression of GTP-Rhoa, Csk or Src pTyr^416^ in microglia (gated using CD45^mid^CD11b^+^) from 4-month-old WT and APP/PS1 mice (n=3-4 mice). **G**, CHME3 microglia expressing a RhoA FRET activity biosensor were recorded in saline before (t=0 min) and after treatment with 200 nM Aβ_1-42_ oligomers (n=20 cells from 3 cultures). **H**, CHME3 microglia expressing a Src FRET activity biosensor were mock-transfected or transfected with RhoA^Q63L^ and then recorded in saline before (t=0 min) and after treatment with 200 nM Aβ_1-42_ oligomers (n=15-16 cells from 3 cultures). **I and J**, CHME3 microglia expressing a NFkB pathway activity biosensor (I) or a glutamate release FRET biosensor (J) were mock-transfected (n=23-25 from 3 cultures) or transfected with RhoA^Q63L^ (n=39 cells from 3 cultures) or incubated with AZD 0530 (200 nM; 1 h; n=31-34 cells from 3 cultures) or pre-incubated with FcR blocker (20 min) followed by treatment with Adalimumab (anti-Tnf; 5 μg/ml; 48 h; n=8 cells from 2 cultures) and then recorded in saline before (t=0 min) and after treatment with 200 nM Aβ_1-42_ oligomers. Data are represented as mean with 95% CI. *P<0.05 (unpaired t test in A-C, F; One-way ANOVA in E; Two-Way ANOVA in G-J). See also Figure S6.

Production of A β by neurons is in large part activity-dependent (Cirrito et al., 2005) and induced by activation of glutamate receptors (Lesne et al., 2005). As decreasing Rhoa activity in microglia enhanced glutamate secretion, we investigated whether this increase modulated the production of Aβ by neurons. The CM from microglia overexpressing the dominant negative Rhoa mutant Rhoa^T19N^ induced a significant increase of Aβ42 in primary hippocampal neurons (**Fig. 5E**). This effect was abrogated by decreasing microglial Src activity or by neutralizing microglial Tnf (**Fig. 5E**).

### Aβ oligomers induce microglia neurotoxic polarization by decreasing Rhoa activity

Our results in Rhoa^fl/fl^:Cx3cr1^CreER+^ mice suggest that microglia dysregulation drives amyloidogenic processing of APP leading to increased production of Aβ by neurons, but could A β signal back to microglia and alter Rhoa activity? Using APP/PS1 mice (Borchelt et al., 1997), in which Aβ is abundantly produced and secreted by neurons, we found a significant decrease of Rhoa activation (GTP-Rhoa) and of Csk expression and an increase of Src activation in APP/PS1 microglia (**Fig. 5F**). No significant differences were found in the numbers of microglia, brain macrophages and blood monocytes (**Suppl. Fig. 6A and B**) or in the percentages of brain macrophages and blood monocytes expressing GTP-Rhoa, Csk or Src Tyr^416^ phosphorylation (**Suppl. Fig. 6C and D**) between 4 mo APP/PS1 and WT littermates.

Next, we investigated whether application of low amounts (200 nM) of Aβ_1-42_ oligomers could indeed modulate Rhoa signaling in microglia. We found by FRET, using the Raichu-Rhoa biosensor (Yoshizaki et al., 2003), that Rhoa activity in living microglia decreased following exposure to Aβ_1-42_ oligomers (**Fig. 5G**). Conversely, Src activity at the plasma membrane (visualized by FRET with the KRas Src YPet biosensor) increased (**Fig. 5H**), and expression of the constitutively active Rhoa mutant Rhoa^Q63L^ significantly attenuated this effect (**Fig. 5H**).

Using the pIκBα-miRFP703 biosensor to detect the activation of canonical NFκB pathway (Shcherbakova et al., 2016), we also observed that exposure to Aβ_1-42_ oligomers induced time-dependent activation of the NFκB pathway in living microglia (**Fig. 5I**), and overexpressing Rhoa^Q63L^ or blocking Src with AZD 0530 abrogated this effect (**Fig. 5I**). Finally, using the FLIPE biosensor we observed by FRET that exposure to Aβ_1-42_ oligomers increased glutamate release from living microglia in a time-dependent manner (**Fig. 5J**), an effect inhibited by Rhoa^Q63L^, by AZD 0530 and by neutralizing Tnf with Adalimumab (**Fig. 5J**). These data suggest that Aβ oligomers can signal back to microglia and decrease RhoA activation, providing a positive feedback loop for Src hyperactivation and Tnf-dependent glutamate release.

### Blocking Src decreases amyloid-like pathology in microglia Rhoa deficient mice

Next, we investigated if inhibiting Src would modulate the reactivity of microglia in early-deposing APP/PS1 mice. Of note, a single IP injection of AZD 0530 in APP/PS1 mice was sufficient to decrease Src activity in their brains (**Suppl. Fig. 6E**). Therefore, and as we did for Rhoa^fl/fl^:Cx3cr1^CreER+^ mice, we blocked Src activity by giving AZD 0530 weekly to 3 mo APP/PS1 mice for 1 month (**Suppl. Fig. 6F**) and investigated whether Src inhibition would preserve hippocampal synapses. Injecting AZD 0530 in APP/PS1 mice substantially inhibited the decrease of vGlut-1^+^/PSD-95^+^ synapses (**Suppl. Fig. 6G**). Concurrent with the idea that Src inhibition decreases microglia activation, we found increased microglial ramification in the neocortex of APP/PS1 mice injected with AZD 0530 (**Suppl. Fig. 6H**). Paralleling this increased microglial ramification in APP/PS1 mice treated with AZD 0530, we also observed an increase in the ramification of microglia following Src inhibition with AZD 0530 in Rhoa^fl/fl^:Cx3cr1^CreER+^ mice (**Suppl. Fig. 6I**).

Plaque-associated microglia restrict amyloid plaque formation limiting pathology in AD mouse models by phagocytosis of amyloid deposits (Wang et al., 2015). We thus investigated whether Rhoa activation and consequent blockade of Src would improve microglial engulfment of Aβ. Chemogenetic activation of Rhoa using the Rhoa-CID strategy significantly increased the uptake of Aβ_1-42_ by microglia (**Fig. 6A**). Activation of Rhoa led to Src inhibition and, accordingly, blockade of Src with AZD 0530 also increased Aβ_1-42_ uptake by microglia (**Fig. 6B**). Blocking Src with AZD 0530 significantly increased microglia engulfment of Aβ deposits in brains of both APP/PS1 (**Suppl. Fig. 6J**) and Rhoa^fl/fl^:Cx3cr1^CreER+^ mice (**Fig. 6C and Suppl. Fig. 6K**). AZD 0530 treatment caused no significant alterations in the amounts of APP either in APP/PS1 or in Rhoa^fl/fl^:Cx3cr1^CreER+^ mice (**Suppl. Fig. 6L and M**), indicating that AZD 0530 does not impact APP production.

**Figure 6.**
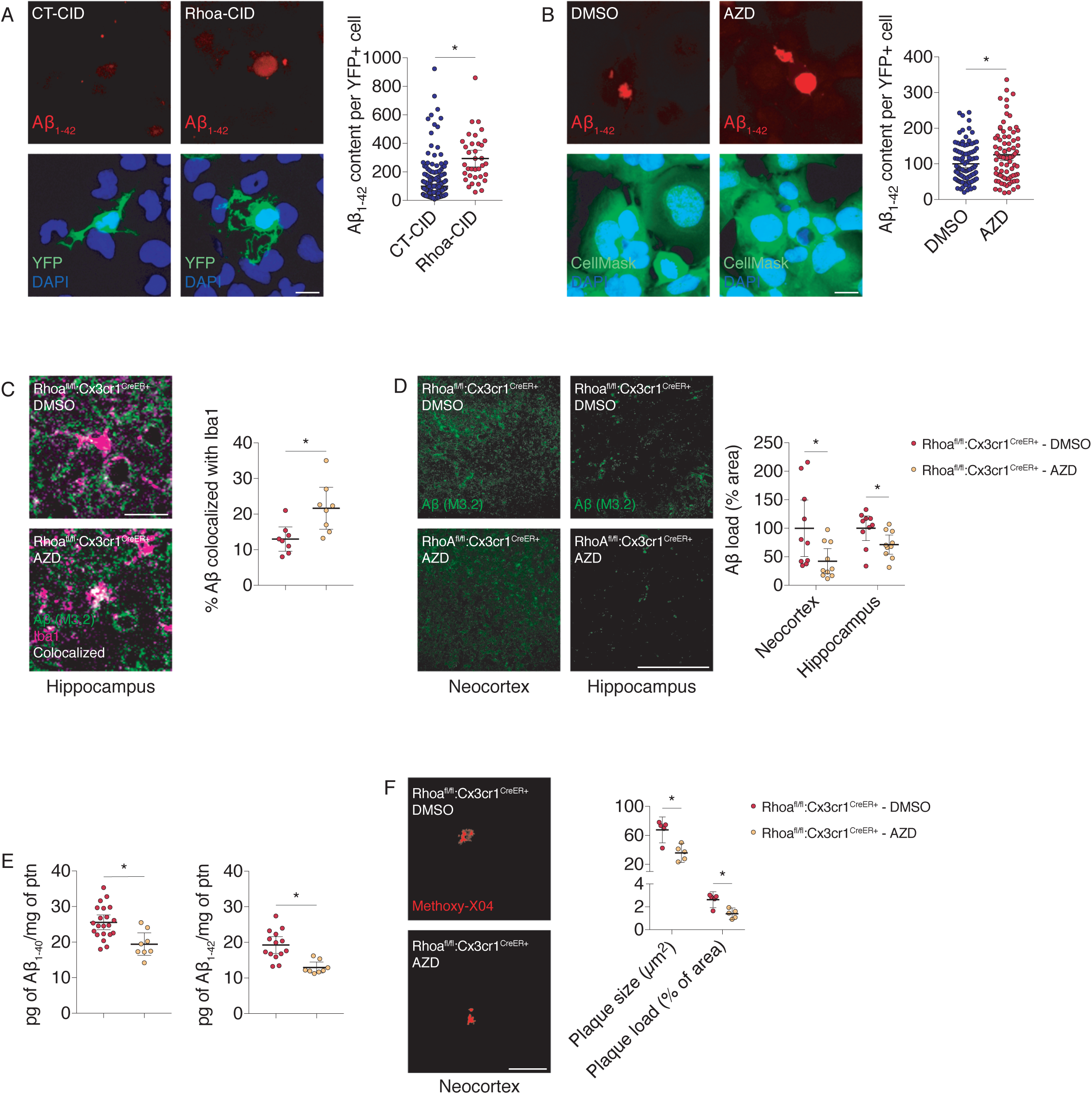
Src inhibition mitigates the amyloid-like pathology elicited by the loss of Rhoa in adult microglia. **A,** CHME3 microglia were co-transfected with YFP-FKBP and Lyn_11_-FRB or with YFP-FKBP-Rhoa (CA) and Lyn_11_-FRB (n=211-395 cells from 3 cultures). All cells were treated with Aβ_1-42_ fluorescent particles (10 µg/ml) and rapamycin (500 nM) for 90 min. **B,** CHME3 microglia (stained with CellMask dye; green) were treated with or AZD 0530 (n=89-113 cells from 3 cultures) and incubated with Aβ_1-42_ fluorescent particles (10 µg/ml) for 90 min. **C**, confocal images showing colocalization (white) of Iba1^+^ microglia with M3.2 immunoreactive amyloid deposits on tissue sections from Rhoa^fl/fl^:Cx3cr1^CreER+^ mice (hippocampus) injected with 10 mg/kg AZD 0530 or DMSO (n=8 mice). Scale bars: 10 μm. **D**, confocal images showing M3.2^+^ amyloid deposits on tissue sections from Rhoa^fl/fl^:Cx3cr1^CreER+^ mice injected with 10 mg/kg AZD 0530 or DMSO (n=10 animals mice). Scale bars: 200 μm. **E**, ELISA for Aβ_1-40_ and Aβ_1-42_ on neocortical extracts of RhoA^fl/fl^:Cx3cr1^CreER+^ mice injected with 10 mg/kg AZD 0530 or DMSO (n=8-21 animals for Aβ_1-40_ and n=8-14 animals for Aβ_1-42_). **F**, confocal images of methoxy-X04^+^ plaque-like deposits on neocortices RhoA^fl/fl^:Cx3cr1^CreER+^ mice injected with 10 mg/kg AZD 0530 or DMSO (n=5 mice). Scale bar: 50 μm. Data are represented as mean with 95% CI. *P<0.05 (Mann-Whitney test in A-D, F; unpaired t test in E). See also Figure S6.

Paralleling the increased Aβ clearance by microglia, we observed a significant decrease of amyloid burden (Aβ amounts detected using BAM-10 and E610 antibodies) in the brain of APP/PS1 mice injected with AZD 0530 (**Suppl. Fig. 6N and O**). AZD 0530 also decreased Aβ deposition (**Fig. 6D and Suppl. Fig. 6P**), the production of both Aβ_1-40_ and Aβ_1-42_ peptides (**Fig. 6E**) and the formation of fibrillar plaque-like structures labeled by methoxy-X04 (**Fig. 6F**) in the neocortex of Rhoa^fl/fl^:Cx3cr1^CreER+^ mice, indicating that blockade of Src downstream of Rhoa mitigates amyloid burden.

## Discussion

Crossing Cx3cr1^CreER-EYFP^ mice (Parkhurst et al., 2013) with Rhoa floxed mice (Herzog et al., 2011, Jackson et al., 2011) allowed studying the role of Rhoa in adult brain microglia and circumvented some limitations of previous studies using overexpression of Rhoa dominant mutants (Bianchi et al., 2011, Ohsawa et al., 2000, Moon et al., 2013), which may trigger off-target and trans-dominant effects (Zhou and Zheng, 2013, Wang and Zheng, 2007), as well as the use of C3 exoenzyme toxin (Rattan et al., 2003, Hoffmann et al., 2008), which not only inhibits Rhoa but also Rhob and Rhoc (Mohr et al., 1992, Chardin et al., 1989).

Rhoa works through multiple downstream pathways and effectors, and it is to be expected that the loss of Rhoa might disturb the functions and/or interactions of more than one of its effectors. For example, knocking out Rhoa in the mouse spinal cord neuroepithelium does not cause cytokinesis defects (Herzog et al., 2011), whereas knocking out the Rhoa effector citron-kinase impairs cytokinesis (Di Cunto et al., 2000). This could also explain why Rhoa deficiency activates microglia whereas the pharmacological inhibition of the Rhoa effector Rho-kinase (ROCK) does not (Mueller et al., 2005, Chen et al., 2013). Another factor potentially contributing for the disparity of these 2 phenotypes is that in addition of being activated by Rhoa, ROCK can also be regulated by Rhoe/Rnd3 (Riento et al., 2003) or by Rhoc (Riento and Ridley, 2003). Our data, however, suggested that Rhoc did not play a significant role in microglia proinflammatory activation and microglia-induced neurotoxicity.

ROCK inhibition by Y27632 or fasudil attenuates microglia activation, promoting some functional recovery in animal models for different neurological conditions (Mueller et al., 2005, Chen et al., 2013, Chong et al., 2017, Sellers et al., 2018). Because Rhoa is likely to exert its control over microglia immune activity through several other downstream effectors besides ROCK, which activation can also be regulated independently of Rhoa (Julian and Olson, 2014), these studies are not very informative concerning Rhoa regulation of microglial function. In addition, Y27632 and fasudil are not cell specific and have substantial off-target effects (Vigil and Der, 2013), making it difficult to attribute any phenotype solely to the inhibition of ROCK downstream of Rhoa.

Excessive secretion of glutamate by microglia can induce excitotoxic neuronal damage (Maezawa and Jin, 2010, Takeuchi et al., 2006), and by exacerbating neuronal activity can also increase the production of Aβ species by neurons (Lesne et al., 2005). Blocking Tnf, and the consequent release of glutamate, from microglia lacking Rhoa prevented neurite damage and also the increase of Aβ42 in primary hippocampal neurons. Injecting pomalidomide (a brain penetrant immunomodulatory agent that also inhibits Tnf expression) in Rhoa mutant mice decreased brain Tnf expression and attenuated synapse loss. Although pomalidomide might have inhibited other pathways in addition to Tnf, our in vitro data demonstrated that Tnf signaling was required for neurite beading and synapse loss, suggesting that Tnf contributed for the synapse loss following Rhoa ablation in microglia.

AZD 0530 inhibits SFKs phosphorylation, including Fyn, Src, Lyn and other family members. Although AZD 0530 inhibits Src more potently than other SFKs (for example the reported IC_50_ for Src is 2.7 nM, while for Fyn is around 10 nM (Green et al., 2009)), and despite of our compelling in vitro results pointing out for a key and rather specific role of Src in driving the phenotype produced by depleting Rhoa in microglia, we cannot claim that, in vivo, the only kinase inhibited was Src. This is also applicable to the work of Kaufmann and colleagues, who report that the effect of AZD 0530 in diminishing some aspects of AD-like pathology in APP/PS1 mice is due to decreased Fyn activation in neurons (Kaufman et al., 2015). It is likely that besides inhibiting Fyn in neurons, AZD 0530 might also inhibit other SFKs in neurons and in other brain cell types, including Src in microglia. Accordingly, and in the context of our work, we cannot therefore exclude that the effect of AZD 0530 in preventing synapse loss and decreasing the amyloid burden in Rhoa mutants and in APP/PS1 mice was contributed, at least in part, by SFK inhibition in both glia and neurons.

Dose-response curve of AZD 0530 revealed an apparent IC_50_ of 0.024 mg/kg for Src inhibition in the brain. AZD 0530 has a half-life of 16 h in the brain (Kaufman et al., 2015), therefore, around 0.0146 mg/kg (10.5 half-lives) of AZD 0530 will remain in the brain following our administration protocol. Because the lowest dose of AZD 0530 (0.002 mg/kg) we tested produced around 30% of Src inhibition, these data show that the dosage of AZD 0530 we used effectively sustained Src inhibition in the brain, which is in accordance with a persistent effect of AZD 0530 reported for APP/PS1 mice after drug washout (Smith et al., 2018).

We observed decreased Rhoa activity in microglia obtained from early-depositing APP/PS1 mice, a stage in which synapse loss was already present without robust amyloid pathology, suggesting that such decrease in Rhoa activation might have contributed to the synapse loss in early-depositing APP/PS1 mice. Indeed, blocking Src, downstream of Rhoa, prevented synapse loss, increased microglial engulfment of Aβ and decreased Aβ burden. However, substantial glial activation driven by overexpressing proinflammatory cytokines can also decrease amyloid burden in AD mice (Chakrabarty et al., 2011, Chakrabarty et al., 2010), casting doubts whether attenuating microglial activation *per se* would be sufficient to preserve synapses as amyloid pathology builds up during the course of AD.

Loss of synapses, LTP impairment, recognition memory deficits and deposition of Aβ are consistent hallmarks found in human AD patients and in animal models of AD. Similarly, Rhoa ablation in adult microglia reproduced some of these hallmarks, further supporting the hypothesis that disrupting Rhoa signaling produces spontaneous microglia activation leading to neurodegeneration. In this regard, our data provide additional validation to notion of “microgliopathy” (Prinz and Priller, 2014), in which microglia dysfunction, by itself, could initiate or be a major cause of neurological disease.

## Acknowledgements

The authors acknowledge the support of the following i3S Scientific Platforms: Animal Facility, Translational Cytometry Unit (TraCy), BioSciences Screening (BS) and Advanced Light Microscopy (ALM), members of the national infrastructure PPBI-Portuguese Platform of BioImaging (supported by POCI-01–0145-FEDER-022122).

FCT Portugal (PTDC/MED-NEU/31318/2017-031318) supported work in JBR lab.

FCT Portugal, PEst (UID/NEU/04539/2013), COMPETE-FEDER (POCI-01-0145-FEDER-007440), Centro 2020 Regional Operational Programme (CENTRO-01-0145-FEDER-000008: BrainHealth 2020) and Strategic Project UIDB/04539/2020 and UIDP/04539/2020 (CIBB) supported work in AFA lab.

CCP and RS hold employment contracts financed by national funds through FCT – Fundação para a Ciência e a Tecnologia, I.P., in the context of the program-contract described in paragraphs 4, 5 and 6 of art. 23 of Law no. 57/2016, of August 29^th^, as amended by Law no. 57/2017 of July 19^th^.

## Author contribution

**Conceptualization**, RS, CCP and JBR; **Methodology**, RS, CCP; **Investigation**, RS, CCP, TC, AR, TOA, JFH, SHV, JM, CMS, FIB, RLA, VSC, APS and AM; **Writing – Original Draft**, RS, CCP, APS, RPC, CB, AMS, TS, AFA and JBR; **Writing – Review & Editing**, RS, CCP and JBR; **Funding Acquisition**, AFA, JBR; **Supervision**, JBR.

## STAR Methods

### RESOURCE AVAILABILITY

#### Lead Contact

Further information and requests for resources and reagents should be directed to and will be fulfilled by the Lead Contact, João Relvas.

#### Resource Availability

This study did not generate new unique reagents.

#### Data and Code Availability

This study did not generate any unique codes.

### EXPERIMENTAL MODEL AND SUBJECT DETAILS

#### Animals

All mice experiments were reviewed by i3S animal ethical committee and were approved by Direção-Geral de Alimentação e Veterinária (DGAV). Animals were maintained in standard laboratory conditions with inverted 12h/12h light dark cycle and were allowed free access to food and water. Animals were group-housed under specific pathogen-free conditions. Experiments were carried out in adult mice (2-3 months of age) following the 3Rs ethics.

##### Conditional Rhoa-deficient mice

All mice experiments were approved by Direção-Geral de Alimentação e Veterinária (DGAV). Cx3cr1^CreER-EYFP^ mice were purchased from Jackson Laboratories. In such mice, the *Cx3cr1* promoter drives high expression of the CreER cassette in microglia (Parkhurst et al., 2013). Mice homozygous for the *Rhoa* floxed allele (Herzog et al., 2011, Jackson et al., 2011) were backcrossed at least for 10 generations and were kept in C57BL/6 background at the I3S animal facility. All genotypes were determined by PCR on genomic DNA. Primers used for Rhoa floxed alleles were: AGC CAG CCT CTT GAC CGA TTT A (forward); TGT GGG ATA CCG TTT GAG CAT (reverse). Primers for CreER insertion were: AAG ACT CAC GTG GAC CTG CT (WT forward); AGG ATG TTG ACT TCC GAG TG (WT reverse); CGG TTA TTC AAC TTG CAC CA (mutant reverse).

Rhoa floxed mice were crossed with Cx3cr1^CreER-EYFP^ mice. Progeny of interest were: Control (Rhoa^fl/fl^) and mutants (Rhoa^fl/fl^:Cx3cr1^CreER+^). Mice were given tamoxifen (10 mg *per* animal by oral gavage) at P26 and P28 and then analyzed at different time points (specified on the figure legends). All experiments were performed on mice kept on a C57Bl/6 background. For electrophysiological recordings and in behavioral tests only male Rhoa^fl/fl^, Cx3cr1^CreER+^ and Rhoa^fl/fl^:Cx3cr1^CreER+^ littermates were used. No differences in the mRNA expression levels of Rhoa were found in FACS-sorted microglia between the brains of Rhoa^fl/fl -^ and wild-type mice after TAM administration. In addition, after TAM administration, Rhoa^fl/fl^ mice displayed no overt difference in the behavioral performance on motor, anxiety-related and memory tests when compared to wild-type mice. Administration of AZD 0530 (used at a final concentration of 10 mg/kg via IP injections; 1 IP per week lasting 4 weeks) to Rhoa^fl/fl^:Cx3cr1^CreER+^ and Rhoa^fl/fl^ mice started 1 week after the second TAM pulse. Administration of pomalidomide (used at a final concentration of 50 mg/kg via IP injections; 3 IPs per week lasting 4 weeks) to Rhoa^fl/fl^:Cx3cr1^CreER+^ and Cx3cr1^CreER+^ mice started 1 week after the second TAM pulse.

##### Tnf deficient mice

Prof. Rui Applelberg (University of Porto) supplied C57BL/6.Tnf knockout (referred herein as Tnf KO) mice. Tnf KO mice were genotyped by PCR using ATC CGC GAC GTG GAA CTG GCA GAA (forward) and CTG CCC GGA CTC CGC AAA GTC TAA (reverse) primer pair. Tnf KO mice display a single band of 2 kb in the PCR gel. Mice were bred at i3S animal facility. Tnf KO animals were bred to maintain maximal heterozygosity.

##### AD mice

The AβPPswe/PS1A246E (termed APP/PS1) AD mice model co-expresses a chimeric mouse-human amyloid-β protein precursor (AβPP) bearing a human Aβ domain with mutations (K595N and M596L) linked to Swedish familial AD pedigrees and human presenilin-1 A246E mutation, with both transgenes under the control of the mouse prion protein promoter (Borchelt et al., 1997). This AD mouse model displays age-dependent and region-specific amyloidosis, gliosis and associated hippocampal-dependent memory deficits, resembling the AD pathology found in human patients (Borchelt et al., 1997). APP/PS1 mice were genotyped by PCR using PSEN primers: AAT AGA GAA CGG CAG GAG CA (forward) and GCC ATG AGG GCA CTA ATC AT (reverse); APP primers: GAC TGA CCA CTC GAC CAG GTT CTG (forward) and CTT GTA AGT TGG ATT CTC ATA TCC G (reverse); WT primers: CCT CTT TGT GAC TAT GGT GAC TGA TGT CGG (forward) and GTG GAT AAC CCC TCC CCC AGC CTA GAC C (reverse). APP/PS1 mice were maintained in SV129 background and were bred at the i3S animal facility. Administration of AZD 0530 (used at a final concentration of 20 mg/kg via IP injections; 1 IP per week lasting 4 weeks) to APP/PS1 mice started when animals completed 3 months of age.

#### Primary cultures of hippocampal neurons

Hippocampi were dissected from embryonic day 18 Wistar rat embryos and dissociated using trypsin (0.25%, v/v). Neurons were plated at a final density of 1×10^4^ to 5×10^4^ cells per dish and cultured in the presence of a glial feeder layer. Cells were cultured in Neurobasal medium supplemented with B27 supplement (1:50, v/v), 25 mM glutamate, 0.5 mM glutamine, and gentamicin (0.12 mg/ml). To prevent glial overgrowth, neuronal cultures were treated with 5 mM cytosine arabinoside after 3 days in vitro (DIV) and maintained in a humidified incubator with 5% CO_2_/95% air at 37°C for up to 2 weeks, feeding the cells once per week by replacing one-third of the medium. To assess neurite beading, hippocampal neurons expressing mVenus were observed under a fluorescence microscope after 24 h of incubation with 50% (v/v) microglia conditioned media. Neurons were observed blindly in three independent hippocampal cultures. The ratio of bead-bearing mVenus^+^ neurons was calculated over the total mVenus^+^ neurons counted.

#### Primary cultures of cortical microglia

Primary cortical microglial cell cultures were performed as previously described (Socodato et al., 2015a, Socodato et al., 2015b, Portugal et al., 2017). In brief, mice pups (2-day-old) were sacrificed, their cerebral cortices were dissected in HBSS, pH 7.2, and digested with 0.07% trypsin plus 50 μL (w/v) DNAse for 15 min. Next, cells were gently dissociated using a glass pipette in DMEM F12 GlutaMAX™-I supplemented with 10% FBS, 0.1% gentamicin. Cells were plated in poly-D-lysine-coated T-flasks (75 cm^2^) at 1.5×10^6^ cells per cm^2^. Cultures were kept at 37°C and 95% air/5% CO_2_ in a humidified incubator. Culture media was changed every 3 days up to 21 days. To obtain purified microglial cell cultures, culture flasks were subjected to orbital shaking at 200 rpm for 2 h. Next, culture supernatant was collected to tubes, centrifuged at 453 g for 5 min at room temperature. The supernatant was discarded and the pellet, containing microglia, was re-suspended in culture medium, and cells were seeded in poly-D-lysine-coated 6 or 12-well culture plates at 2.5×10^5^ cells/cm^2^ with DMEM F12 GlutaMAX™-I supplemented with 10% FBS, 0.1% gentamicin and 1 ng/ml M-CSF or 1 ng/ml GM-CSF. Purified microglia were cultured for 5-8 days. Immunolabeling with CD11b showed a purity of 95-99% for these cultures.

#### Microglial cell lines

The microglial cell line N9 was obtained by immortalization of primary cultures from the ventral mesencephalon and cerebral cortex from ED12-13 CD1 mouse embryos with the 3RV retrovirus carrying an activated v-myc oncogene (Righi et al., 1989). Cells were cultured in RPMI 1640 supplemented with 5% FBS, 23.8mM sodium bicarbonate, 30mM D-Glucose, 100 U/mL penicillin and 100 μg/mL streptomycin, and were maintained at 37°C, 95% air and 5% CO_2_ in a humidified incubator.

The microglial cell line CHME3 was obtained from primary cultures of human embryonic microglial cells by transfection with a plasmid encoding for the large T antigen of SV40 (Janabi et al., 1995). These cells were cultivated with DMEM GlutaMAX™-I supplemented with 10% FBS and 100 U/mL penicillin and 100 μ/mL streptomycin, and were maintained at 37°C, 95% air and 5% CO_2_ in a humidified incubator.

For generation of microglial stable cell sub-clones, N9 or CHME3 cultures were infected with viral particles and allowed to grow for additional 48h. Cultures were then selected using puromycin as before (Socodato et al., 2015b, Portugal et al., 2017). Stable cell sub-clones carrying an empty vector (pLKO) or the insert of interest were validated by Western blotting or by qRT-PCR.

Primary microglial cells, N9 and CHME3 microglial cell lines were transfected using jetPRIME^®^ (Polyplus Transfection) according to the manufacturer’s protocol.

### METHOD DETAILS

#### Flow cytometry and cell sorting

For characterization of microglia and macrophages in the samples, the following markers were used: CD45-PE, CD11b-APC, CD11b-Alexa Fluor 647, CCR2-PE/Cy7 and Ly6C-PerCP/Cy5.5. Microglial and macrophages were collected from the tissues (brain, blood and spleen) of control and mutant mice using density gradient separation. In brief, mice were anesthetized and then perfused with ice-cold PBS. For single cell suspensions, tissues were quickly dissected, placed on ice-cold RPMI and mechanically homogenized. Cell suspension was passed through a 100 μm cell strainer and centrifuged over a discontinuous 70%/30% Percoll gradient. Cells on the interface were collected, pelleted, washed extensively and then counted in a Neubauer chamber using trypan blue exclusion to estimate the number of live cells. Single cell suspensions (5 × 10^5^ cells) were incubated with different mixes of FACS antibodies for 30 min at 4°C in the dark. Compensation settings were determined using spleen from both control and mutant. Cell suspensions were evaluated on a FACS Canto II analyzer (BD Immunocytometry Systems). For intracellular staining, cell suspensions were run over Percoll gradient, washed and counted as above. Afterwards, cells were seeded in a U bottom 96-well plate and incubated with CD45-PE, CD11b-Alexa Fluor 647 or CD11b-PE/Cy7. After antibody washing, cells were fixed in 2% PFA for 30 min, washed in PBS and permeabilized with permeabilization buffer (Life Technologies 00-8333-56). Intracellular antibody staining mix was prepared in permeabilization buffer and incubated with the cells overnight at 4°C in the dark. After washing in permeabilization buffer, cells were incubated with Alexa Fluor 488 or 647 secondary antibody for 1 h at 25°C in the dark. After that, cells were washed twice in permeabilization buffer, washed twice in FACS staining buffer (2% BSA, 0.1% sodium azide in PBS) and analyzed in FACS Canto II. Fluorescence minus one controls for Rhoa, GTP-Rhoa, Csk and phospho-Src were prepared in mixes with CD45-PE and CD11b-Alexa Fluor 647 CD11b-PE/Cy7 plus secondary antibodies alone. Cell sorting was performed on a FACS ARIA cell sorter. Data were analyzed by FlowJo X10 software (TreeStar).

#### MACS isolation of adult microglia

Mice were sacrificed in a CO_2_ chamber and the brains were removed. The right hemisphere was mechanically dissociated in ice-cold Dounce buffer (15mM HEPES; 0,5% Glucose; and DNAse in PBS) by 6 strokes in a tissue potter. Then, the homogenate was passed through a 70 μm cell strainer, centrifuged and cells were counted in a TC10 Automated Cell Counter from BIO-RAD using Trypan Blue to exclude dead cells. 4 × 10^7^ cells were pelleted by centrifugation (9300 g; 1 min; 4°C) and resuspended in 360 μL of MACS Buffer (0.5% BSA; 2 mM EDTA in PBS) followed by incubation with 40 μL CD11b Microbeads (130-093-634 Miltenyi Biotec). CD11b^+^ fraction was selected using LS columns (130-042-401 Miltenyi Biotec) in a MACS separator (Miltenyi Biotec) according to the manufacture’s instructions. Eluted CD11b enriched fraction was centrifuged (9300 g; 1 min; 4°C), the pellet was lysed in RIPA-DTT buffer and used for protein isolation.

#### Brain tissue preparation and immunofluorescence

After animal perfusion with ice-cold PBS (15 ml) and fixation by perfusion with 4% PFA, brains were post-fixed by immersion in 4% PFA in PBS, pH 7.4 overnight. After that, brains were washed with PBS and then cryoprotected using sucrose gradient in a row (15 and 30%). After 24 h, brains were mounted in OCT embedding medium, frozen and cryosectioned in the CM3050S Cryostat (Leica Biosystems). Coronal sections from brains (30 μm thickness between bregma positions 1.0 mm and −2.0 mm) were collected non-sequentially on Superfrost ultra plus slides. Brain sections from control and from mutant mice encompassing identical stereological regions were collected on the same glass slide. Slides were stored at −20°C until processed for immunohistochemistry.

Frozen sections were defrosted by at least 1 h and hydrated with PBS for 15 min. Sections were permeabilized with 0.25% Triton X-100 for 15 min, washed with PBS for 10 min and blocked (5% BSA, 5% FBS, 0.1% Triton X-100) for 1 h. Primary antibodies were incubated in blocking solution in a humidified chamber overnight at 4°C. Secondary antibodies were incubated for 2 h in blocking solution. After the secondary antibody, sections were washed three times for 10 min with PBS and incubated for 10 min with DAPI and rinsed twice in PBS. Slides were cover slipped using glycergel or Immumount and visualized under a Leica TCS SP5 II confocal microscope.

#### Image reconstruction and morphometric analysis

Images from tissue sections were acquired using a Leica HC PL APO Lbl. Blue 20x /0.70 IMM/CORR or a Leica HC PL APO CS 40x /1.10 CORR water objective in 8-bit sequential mode using standard TCS mode at 400 Hz and the pinhole was kept at 1 airy in the Leica TCS SP5 II confocal microscope. Images were resolved at 1024 × 1024 pixels format illuminated with 2-5% DPSS561 561 nm wave laser using a HyD detector in the BrightR mode and entire Z-series were acquired from mouse brain sections. Equivalent stereological regions were acquired for all tissue sections within a given slide. Image series were deconvolved using the Hyugens Professional using the Classic Maximum Likelihood Estimation (CMLE) algorithm together with a determined theoretical PSF established using a routine-based implementation for the Hyugens software. Reconstruction and generation of 3D volumes of deconvolved images were performed using the ImageJ 3D viewer plugin and cell counts were performed blinded on integral 3D volume-rendered images. Sholl analyses were performed using an ImageJ Sholl plugin (Ferreira et al., 2014).

All immunostaining in sets of slides were performed together, using the same batch of primary and secondary antibodies, and blocking and washing solutions. Furthermore, images from different sections within a given slide were acquired on the same day, always by the same operator and with identical microscope parameters (e.g., same power for the confocal Argon laser; same laser-line potency; same objective; same fluorescence exposure times and offset for a given fluorophore; same camera binning, zoom and ROI magnification; same pinhole aperture; same pixel size; same line averaging; same TCS scanner mode and speed; same z-stack step size and optical sectioning).

##### NeuN quantification

Number of NeuN^+^ neurons were manually scored from stereological identical images (3D volume-rendered) of the CA1 region of the hippocampus of stained sections (6 images per hippocampal section; 6 hippocampal sections per animal).

##### Quantification of amyloid deposits

Briefly, M3.2, BAM-10 or 6E10 immunostained sections covering layers II/III and IV of the neocortex or CA1 region of the hippocampus (4 images per section; 6 sections (BAM-10 and 6E10) or 10 sections (M3.2) per animal) were imaged, converted into 8-bit gray scale, 3D volume-rendered and thresholded to highlight immunostained amyloid deposits. Using FIJI software, the percent of immunostained area was calculated for each field and each section.

##### APP quantification

Briefly, APP immunostained sections covering layers II/III and IV of the neocortex (4 images per section; 6 sections per animal) were imaged, converted into 16-bit gray scale, and thresholded to highlight immunostained APP amounts. Using FIJI software, the percent of immunostained area was calculated for each field and each section.

##### Methoxy-X04 quantification

Briefly, stained sections covering layers II/III and IV of the neocortex (5 images per section; 4 sections per animal) were imaged, converted into 16-bit gray scale, and thresholded to highlight stained. Using FIJI software, the percent of stained area was calculated for each field and each section.

##### Quantification of synapses

Images from stereological identical hippocampal CA1 regions from each experimental group (3 images per hippocampi; 4 hippocampal sections per animal) were acquired using a Leica HC PL APO CS 40x /1.10 CORR water objective at 1024 × 1024 pixels resolution with 8-bit bidirectional non-sequential scanner mode at 400 Hz and pinhole at 1 airy in the Leica TCS SP5 II confocal microscope. Z-stacks were converted to maximum projection images using LAS AF routine and the LAS AF colocalization plugin processed each projection using subtracted background (25–36% offset for both channels) and thresholded foreground (35-45% offset for vGlut-1 channel; 30-40% offset for PSD-95 channel). Values corresponding to the positive area of vGlut-1/PSD-95 colocalization puncta (synapses) and values for the overall image area for each image was extracted using the LAS AF colocalization plugin and statistically evaluated in GraphPad Prism.

#### TUNEL assay

Slides were defrosted at room temperature for at least 1 h. After that, sections were hydrated in PBS and fixed with 4% PFA for 10 min. After washes with PBS, sections were blocked (10% NGS; 0,1% BSA and 1% Triton X-100) for 1 h and then equilibrated in TdT buffer (30 mM Tris-HCl; 140 mM Sodium Cacodylate; 1 mM Cobalt chloride; pH 7.2) for 10 min at room temperature. To perform the TUNEL reaction was performed using 1.6 µL of TUNEL Enzyme (11767305001; Roche); 1.2 µL of Biotin-16-dUTP (11093070910; Roche) and 1.2 µL of 1 mM dATP in 200 µL of TdT buffer. The reaction was performed at 37°C for 90 min. The reaction was stopped by incubation with 2x SSC buffer for 10 min. After washes with PBS, Cy3-conjugated Streptavidin (1:200 in blocking solution) and DAPI (2 ug/ml) were incubated for 1 h at room temperature. After PBS washes, sections were mounted with fluoroshield (Sigma-Aldrich). The number of TUNEL^+^ cells was manually scored from stereological identical images of the CA1 region of the hippocampus of stained sections (4 images per hippocampal section; 8 hippocampal sections per animal).

#### Quantification of Iba1 colocalization with amyloid deposits

Images from stereological identical neocortical or hippocampal CA1 regions from each experimental group (4 images per section; 6 sections per animal) were acquired using a Leica HC PL APO CS 40x /1.10 CORR water objective at 1024 × 1024 pixels resolution with 8-bit bidirectional scanner mode at 200 Hz in the Leica TCS SP5 II confocal microscope. Z-stacks were converted to maximum projection images using LAS AF routine and the LAS AF colocalization plugin processed each projection using subtracted background (45-55% offset for both channels) and thresholded foreground (65% offset for BAM-10/M3.2 channel; 20-35% offset for Iba1 channel). Values corresponding to the positive area of colocalization puncta and values for the overall image area for each image were extracted using the LAS AF colocalization plugin and statistically evaluated in GraphPad Prism.

#### Immunofluorescence on cultured microglia

Microglial cultures were cultivated in glass coverslips. Coverslips were fixed with 4% PFA, washed three times for 5 min in PBS, permeabilized with 0.1% Triton X-100 for 10 min, washed again and incubated for 1 h in blocking solution (5% BSA). Primary antibodies were added in blocking solution and coverslips were maintained in a humidified chamber for 1 h. Coverslips were washed three times for 10 min with PBS and incubated with secondary antibodies for 1 h in blocking solution. Next, coverslips were washed three times for 10 min with PBS. After the last secondary antibody (when double-labeling was performed), the coverslips were incubated for 1 min with 1 mg/ml DAPI, and rinsed twice in PBS. Coverslips were mounted with Immumount, visualized in a Leica DMI6000B inverted epifluorescence microscope using PlanApo 63X/1.3NA glycerol immersion objective. Images were acquired with 4 × 4 binning using a digital CMOS camera (ORCA-Flash4.0 V2, Hamamatsu Photonics).

For quantification, images were exported as raw 16-bit tiff using the LAS AF software. Background was subtracted in FIJI using the roller-ball ramp in between 35-50% pixel radius. Images were segmented in FIJI using the local Otsu threshold. Thresholded images were converted to binary mask using the dark background function. Binary mask images were multiplied for their respective original channel images using the image calculator plug-in to generate a masked 32-bit float images relative to each channel. Original coordinate vectors were retrieved from the ROI manager and FIJI returned the mean fluorescent intensity in gray values contained within any single microglia using the multi-measure function. Mean fluorescent intensity for each single microglia were exported and statistically evaluated with the GraphPad Prism software.

#### Time-lapse video microscopy and FRET assays

For live cell imaging, microglia were plated on plastic-bottom culture dishes (µ-Dish 35 mm, iBidi). Imaging was performed using a Leica DMI6000B inverted microscope. The excitation light source was a mercury metal halide bulb integrated with an EL6000 light attenuator. High-speed low vibration external filter wheels (equipped with CFP/YFP excitation and emission filters) were mounted on the microscope (Fast Filter Wheels, Leica Microsystems). A 440-520nm dichroic mirror (CG1, Leica Microsystems) and a PlanApo 63X 1.3NA glycerol immersion objective were used for CFP and FRET images. Images were acquired with 2×2 binning using a digital CMOS camera (ORCA-Flash4.0 V2, Hamamatsu Photonics). Shading illumination was online-corrected for CFP and FRET channels using a shading correction routine implemented for the LAS AF software. At each time-point, CFP and FRET images were sequentially acquired using different filter combination (CFP excitation plus CFP emission (CFP channel), and CFP excitation plus YFP emission (FRET channel), respectively).

Quantification of biosensors was performed as before (Portugal et al., 2017, Socodato et al., 2015a, Socodato et al., 2015b, Socodato et al., 2018). In brief, images were exported as 16-bit tiff files and processed in FIJI software. Background was dynamically subtracted from all frames from both channels. Segmentation was achieved on a pixel-by-pixel basis using a modification of the Phansalkar algorithm. After background subtraction and thresholding, binary masks were generated for the CFP and FRET images. Original CFP and FRET images were masked, registered and bleach-corrected. Ratiometric images (CFP/FRET for KRas Src YPet probe and FLIPE glutamate release probe or FRET/CFP for Raichu-Rhoa and Flare.Rhoc sensors) were generated as 32-bit float-point tiff images. Values corresponding to the mean gray values were generated using the multi calculation function in FIJI and exported as mentioned above. KRas Src and glutamate (pDisplay FLIPE-600n^SURFACE^) biosensors have been previously used in microglia giving reliable FRET to donor signals within the probe dynamic range (Socodato et al., 2015b, Socodato et al., 2015a, Socodato et al., 2018, Portugal et al., 2017).

For quantification of microglial protrusion velocity, HMC3 microglia expressing Lifeact-mCherry were incubated in DMEM without phenol red plus 15 mM HEPES and recorded (5 min frames for 90 min at 37°C) using appropriate dichroics for red fluorescent protein. Time-lapse images were exported as 16-bit tiff files and processed in FIJI software using the multi kymograph plugin. Protrusion velocities from Lifeact kymograms were extracted using a routine macro from the multi kymograph package in ImageJ.

#### Preparation of lysates (tissue and cell cultures)

For ELISA, brains were removed, different brains areas were isolated, fast frozen on dry ice and stored at −80°C. Cortices were defrosted and 400 ul of homogenization buffer (50 mM Tris-Base, 4 mM EDTA, 150 mM NaCl, 0.1% Triton X 100 and protease inhibitor cocktail containing AEBSF (Sigma – P8340)). Samples were sonicated (10 pulses of 1 sec at 60Hz) and centrifuged for 15 min at 2271 g at 4°C. The pellets were discarded and samples were centrifuged again at 13500 g for 5 minutes at 4°C. The supernatant was used for ELISA and the total amount of protein were estimated using the Pierce BCA protein assay kit.

For all other applications, cell cultures or mice tissues were lysed using RIPA-DTT buffer (150 mM NaCl, 50 mM Tris, 5 mM EGTA, 1% Triton X-100, 0.5% DOC, 0.1% SDS) supplemented with complete-mini protease inhibitor cocktail tablets, 1 mM DTT and phosphatase inhibitor cocktail. Samples were sonicated (6 pulses of 1 sec at 60Hz) and centrifuged at 16,000 g, 4°C for 10 min. The supernatants were collected and the protein concentration was determined by the Pierce BCA protein assay kit. All samples were denatured with sample buffer (0.5 M Tris-HCl pH 6.8, 30% glycerol, 10% SDS, 0.6 M DTT, 0.02% bromophenol blue) at 95°C for 5 min and stored at −20°C until use.

#### Enzyme-linked immunosorbent assay (ELISA)

The concentration of Tnf, Aβ_1-40_ and Aβ_1-42_ were quantified by ELISA following the instructions provided by the manufacturer (Peprotech, UK). Absorbance at 405 nm, with wavelength correction at 650 nm, was measured with a multimode microplate reader (Synergy HT, Biotek, USA). Values corresponding to pg/ml were obtaining by extrapolating a standard concentration curve using recombinant Tnf.

#### Glutamate determination by fluorimetry

Extracellular glutamate was measured in microglia culture supernatant by the formation of NADH in a reaction catalyzed by glutamate dehydrogenase essentially as elsewhere (Socodato et al., 2015b). Briefly, Tris– acetate (75 mM, pH 8.4) plus NAD and ADP were added to culture supernatants and initial fluorescence arbitrary unit (FAU_i_) was quantified in a Luminometer. Glutamate dehydrogenase was added for 30 min to promote the formation of alpha-ketoglutarate and NADH. NADH fluorescence was also determined (FAU_f_). Relative extracellular glutamate content was plotted as FAU_f_ - FAU_i_.

#### Nuclear and cytosolic fractionation

Cultures were washed twice with PBS, and cells were scraped off culture dishes with lysis buffer [50 mM tris-HCl, 5 mM EDTA, 100 mM NaCl, 1 mM DTT, 100 mM phenylmethylsulfonyl fluoride, 200 mM Na_3_VO_4_, 1% Triton X-100, leupeptin (4 mg/ml)]. Lysates were centrifuged (16,000g for 10 min at 4°C), the supernatant (cytosolic fraction) was collected, and the pellet (nuclear fraction) was resuspended. All samples were denatured with sample buffer for 5 min at 95°C and stored at −20°C. Both fractions were subjected to SDS–polyacrylamide gel electrophoresis (SDS-PAGE). To confirm the efficacy of the separation assay, we used LAMP-1 (Enzo Life Sciences) as a cytosolic fraction marker and histone H3 (Millipore) as a nuclear fraction marker.

#### Western blotting

Samples were separated in SDS-PAGE, transferred to PVDF membranes, which were incubated overnight with primary antibodies. Membranes were washed in TBS-T buffer pH 7.6, incubated with peroxidase-conjugated secondary antibodies and developed using an ECL chemiluminescence kit or an ECF fluorescence kit. Images were acquired in a Typhoon FLA 9000 system or ChemiDoc XRS System (Bio-Rad) and quantified by FIJI software.

#### Electrophysiology

For electrophysiological recordings, acute hippocampal slices from P65-80 male control and Rhoa mutants mice were prepared. All procedures were carried out according to the European Union Guidelines for Animal Care (European Union Council Directive 2010/63/EU) and Portuguese law (DL 113/2013) with the approval of the Institutional Animal Care and Use Committee.

Animals were sacrificed by decapitation after cervical displacement and the brain was rapidly removed in order to isolate the hippocampus. Hippocampi were dissected in ice-cold artificial cerebrospinal fluid (aCSF) containing: 124 mM NaCl, 3 mM KCl, 1.2 mM NaH_2_PO_4_, 25 mM NaHCO_3_, 2 mM CaCl_2_, 1 mM MgSO_4_ and 10 mM glucose), which was continuously gassed with 95%O_2_/5% CO_2_. Hippocampal slices were quickly cut perpendicularly to the long axis of the hippocampus (400 μm thick) with a McIlwain tissue chopper and allowed to recover functionally and energetically for at least 1 h in a resting chamber filled with continuously oxygenated aCSF, at room temperature (22–25°C), before being set up for electrophysiological recordings.

For extracellular recordings of field Excitatory Post-Synaptic Potentials (fEPSP), slices were transferred to a recording chamber for submerged slices (1 ml capacity plus 5 ml dead volume) and were constantly superfused at a flow rate of 3 ml/min with aCSF kept at 32°C, gassed with 95% O_2_ – 5%CO_2_. Evoked fEPSP were recorded extracellularly using a microelectrode (4 to 8 M extracellularly using feraCSF solution placed in the *stratum radiatum* of the CA1 area). fEPSP data were acquired using an Axoclamp-2B amplifier (Axon Instrumnets, USA). fEPSPs were evoked by stimulation through a concentric electrode to the Schaffer collateral fibres. Each individual stimulus consisted of a 0.1 ms rectangular pulse applied once every 20s, except otherwise indicated. Averages of six consecutive responses were continuously acquired, digitized with the WinLTP program (Anderson and Collingridge, 2001) and quantified as the slope of the initial phase of the averaged fEPSPs. The stimulus intensity was adjusted at the beginning of the experiment to obtain a fEPSP slope close to 1 mV/ms, which corresponded to about 50% of the maximal fEPSP slope.

θ-burst stimulation was used to induce LTP because this pattern of stimulation is considered closer to what physiologically occurs in the hippocampus during episodes of learning and memory in living animals (Albensi et al., 2007). After obtaining a stable recording of the fEPSP slope, a θ-burst of stimuli, consisting of 1 train of 4 bursts (200 ms interburst interval), each burst being composed of 4 pulses delivered at 100 Hz [1 × (4 × 4)], was applied, and the stimulus paradigm was then resumed to pre-burst conditions up to the end of the recording period (60 min after burst stimulation). LTP magnitude was quantified as % change in the average slope of the fEPSP taken from 50-60 min after LTP induction as compared with the average slope of the fEPSP measured during the 10 min before induction of LTP. fEPSPs were recorded under basal stimulation conditions (standard stimulus intensity and frequency) and stability of fEPSP slope values guaranteed for more than 10 min before changing any protocol parameter. One or two slices per animal were tested at each experimental day.

#### Behavioral tests

Testing procedures were conducted in the dark phase of the light/dark cycle. Only male mice were used in behavioural analysis. Before each session, mice were removed from their home cage in the colony room and brought into the adjacent testing rooms (illuminated with 100 lux and attenuated noise). Behavioural tests were performed in 3 consecutive days in the following order: (1) elevated plus-maze; (2) open field; (3) novel object recognition. All tests were video-recorded. In the elevated plus-maze and open-field tests movement and location of mice were analyzed by an automated tracking system equipped with an infrared-sensitive camera (Smart Video Tracking Software v 2.5, Panlab, Harvard Apparatus). Data from the object recognition test were analyzed using the Observer 5 software (Noldus Information Technology, Wageningen, The Netherlands). All apparatus were thoroughly cleaned with neutral soap after each test session.

##### Elevated plus-maze (EPM)

The maze was made of opaque grey PVC consisting of four arms arranged in a plus-shaped format; two arms have surrounding walls (closed arms, 37×6 cm x18 cm-high), while the two opposing arms have no walls (open arms, 37×6 cm). The apparatus is elevated by 50 cm above the ground. Mice were placed on the central platform facing an open arm and were allowed to explore the maze for 5 min. Open arms entries and time spent in open arms were obtained automatically (video tracking) and used to assess anxiety-like behavior.

##### Open field (OF)

Mice were placed in the center of an OF apparatus (40 × 40 × 40 cm) and then allowed to move freely for 10 min. The distance travelled, peripheral activity and center activity (locomotion in the central section of the OF) were obtained automatically (video tracking).

##### Novel object recognition (NOR)

The NOR test was performed as previously described (Leger et al., 2013). Briefly, the test apparatus consists of an open box and the objects used are made of plastic, glass or metal in three different shapes: cubes, pyramids and cylinders. The test consists of three phases. During habituation phase mice are allowed to explore the apparatus for 10 min (time used to perform OF test). The following day, the acquisition/sample phase starts by placing each mouse in the apparatus with two identical objects (familiar) for 10 min. Then the mouse goes back to its home cage. After 4 h (inter-trial interval, ITI), the retention/choice session is performed. In this phase, the apparatus contains a novel object and a copy of the previous familiar object; animals are allowed to explore these objects for 3 min. Exploration was defined as follows: mouse touched the object with its nose or the mouse’s nose was directed toward the object at a distance shorter than 2 cm (Ennaceur et al., 2005). Circling or sitting on the object was not considered exploratory behavior. Increased time spent exploring the novel object serves as the measure of recognition memory for the familiar object. Positive difference between the exploration time for novel and familiar objects and a discrimination index (DI) above 0.5 were used as indicators of object recognition (Ennaceur et al., 2005). The discrimination index (DI) was calculated as index of memory function, DI = (time exploring the novel object) / (total time spent exploring both objects) (Ennaceur et al., 2005).

#### Cloning of shRNAs into pSicoR-GFP vector

The cloning was performed as previous described (Ventura et al., 2004). Briefly, shRNA oligos (Rhoa sense: TGTCAAGCATTTCTGTCCTTCAAGAGAATTTGGACAGAAATGCTTGACTTTTTTC and antisense: CGAGAAAAAAGTCAAGCATTTCTGTCCAAATTCTCTTGAAATTTGGACAGAAATGCTTGACA; Rhob sense: TCGACGTCATCCTTATGTGCTTTTCAAGAGAAAGCACATAAGGATGACGTCGTTTTTTC and antisense: TCGAGAAAAAACGACGTCTTATGTGCTTTCTCTTGAAAAGCACATAAGGATGACGTCGA; Rhoc sense: TACCTG AGGCAAGATGAGCATATTCAAGAGATATGCTCATCTTGCCTCAGGTTTTTTTC and antisense: GAGAAAAAAACCTGAGGCAAGATGAGCATATCTCTTGAATATGCTCATCTTGCCTCAGGTA were annealed and cloned into *Hpa*I-*Xho*I-digested pSicoR using the T_4_ ligase. Oligos insertion into the vector was validated using PCR and confirmed by sequencing. A control shRNA with no mammalian target (directed against DsRed2; referred to as scrambled) was used in experiments using the pSicoR as before (Portugal et al., 2017).

#### Lentiviruses/Retroviruses production

This protocol was performed exactly as described elsewhere (Socodato et al., 2012, Mejia-Garcia et al., 2013). Low passage HEK293T cells were seeded in 100 mm culture dishes. When cultures reached ∼80% confluence cells were co-transfected overnight with viruses-producing plasmids using jetPRIME^®^. Transfection ratios were as follows: 6 μg of shRNA plasmids to 3 μg of psPAX2 to 3 μg of VSVG (2:1:1) for lentiviruses production or 8 μg of VSVG (or Src^Y416F^ or Src^Y527F^ construct to 4 μg of pUMVC to 2 μg of VSVG (4:2:1) for retroviruses production. The next day, normal growth media replaced transfection media and cells were cultivated for an additional 48 h. Next, media with viral particles were collected, centrifuged at 906 g for 15 min at 4°C, and the supernatant was collected into new tubes and kept at −80°C.

#### Total RNA extraction, cDNA synthesis and qRT-PCR

Total cerebral cortex RNA was extracted using the Direct-zol™ RNA MiniPrep kit according to the manufacturer’s instructions. RNA from FACS-sorted microglia was harvested using the Quick-RNA™ MicroPrep kit. RNAs quality and concentration were all determined using a NanoDrop ND-1000 Spectrophotometer. cDNA synthesis was performed using 100-500 ng of total RNA (DNase I treated) with SuperScript^®^ III First-Strand Synthesis SuperMix.

qRT-PCR was carried out using iQ™ SYBR^®^ Green Supermix on an iQ™5 multicolor real-time PCR detection system (Bio-Rad). The efficiency was analyzed using a log-based standard curve. Expression of PCR transcripts between genotypes was calculated using the 2^-deltaCt^ method with *Yhwaz* serving as the internal control gene.

#### Preparation of Aβ oligomers

Human Aβ_1-42_ peptide (Genscript) was dissolved to 2 mM in DMSO essentially as before (Silva et al., 2017). For oligomer formation, Aβ_1-42_ peptide was diluted to 100 μM in Ca^2+^-free HBSS and incubated for 24-48 h at 4°C. Oligomers were used at final concentration of 200 nM in microglial cultures.

#### Aβ_1-42_ uptake assay

HMC3 microglial cells were seeded on 96-well plate (PerkinElmer) at 1×10^4^ cells per well using an automated Multidrop dispenser (Thermo Scientific) and incubated for 24 hours at 37°C in 5% CO_2_. Then, cells were co-transfected with FKBP-YFP and FRB-Lyn and or with Rhoa-YF (CA) and FRB-Lyn. For chemogenetic activation of Rhoa, cells were pre-treated with rapamycin (Sigma; 500 nM) for 15 min. Different non-transfected cultures were also pre-incubated with AZD 0530 (Src inhibitor; 500 nM) or vehicle (DMSO) for 15 min. Afterwards, cells were incubated with 10 µg/ml Aβ(1–42) HiLyte™ Fluor 555-labeled (Eurogentec) for 90 min at 37 °C in 5% CO_2_. Cells were then thoroughly washed, fixed with 4% MP-paraformaldehyde (w/v), permeabilized with 0.1% (v/v) Triton X-100 (Sigma) for 30 min at room temperature and labelled with DAPI and HCS CellMask™ (Invitrogen).

Images of microglial cultures were acquired using IN Cell Analyzer 2000 microscope (GE Healthcare) using a large chip CCD Camera (binning 2X2) and a Nikon 20x/0.45 NA Plan Fluor objective. Images of Aβ_1-42_, DAPI-stained nuclei, HCS CellMask™ and in some conditions transfected (YFP^+^) cells were acquired using appropriate dichroics for TexasRed, DAPI, Cy5 and FITC channels, respectively. For Aβ_1-42_ signal quantification, the CellProfiler (3.1.8) software was used. For nuclei segmentation, a global thresholding strategy was used followed by Otsu local thresholding. Clumped microglial nuclei were distinguished and individualized. Afterwards, microglial cells were segmented using the raw HCS CellMask™ image through a propagation algorithm based on the minimum cross entropy thresholding method applied to the HCS CellMask™ channel using segmented nuclei as start point and then spanning up to the edges of the labelled microglial plasma membrane. The mean fluorescent intensity of Aβ_1–42_ HiLyte™ Fluor 555 in each microglial cell was retrieved and plotted for statistical evaluation.

### QUANTIFICATION AND STATISTICAL ANALYSIS

Graphs are displayed as dot plots showing full data dispersion with the mean and the 95% confidence interval of the mean. P<0.05 was the cut-off for considering statistically significant difference between comparisons in sampled groups. Experimental units in biological replicates were evaluated a priori for Gaussian distribution using the D’Agostino & Pearson omnibus normality test. When comparing only 2 experimental groups, the unpaired Student t test with equal variance assumption was used for data with normal distribution, while the Mann-Whitney test was used for with non-parametric data. When comparing 3 or more groups, ordinary One-way ANOVA followed by the Bonferroni’s multiple comparations test was used for data with normal distribution, and the Kruskal-Wallis test followed by Dunn’s multiple comparations test was used for non-parametric data. When comparing 4 groups with 2 independent variables, Two-way ANOVA followed by the Bonferroni’s multiple comparations test was used. All imaging quantifications (cell cultures, FACS or mice tissue sections) were blind performed. All statistical analyses were carried out using the Graph Pad Prism 7.0 software. Further statistical details are indicated in the figure legends.

### KEY RESOURCES TABLE

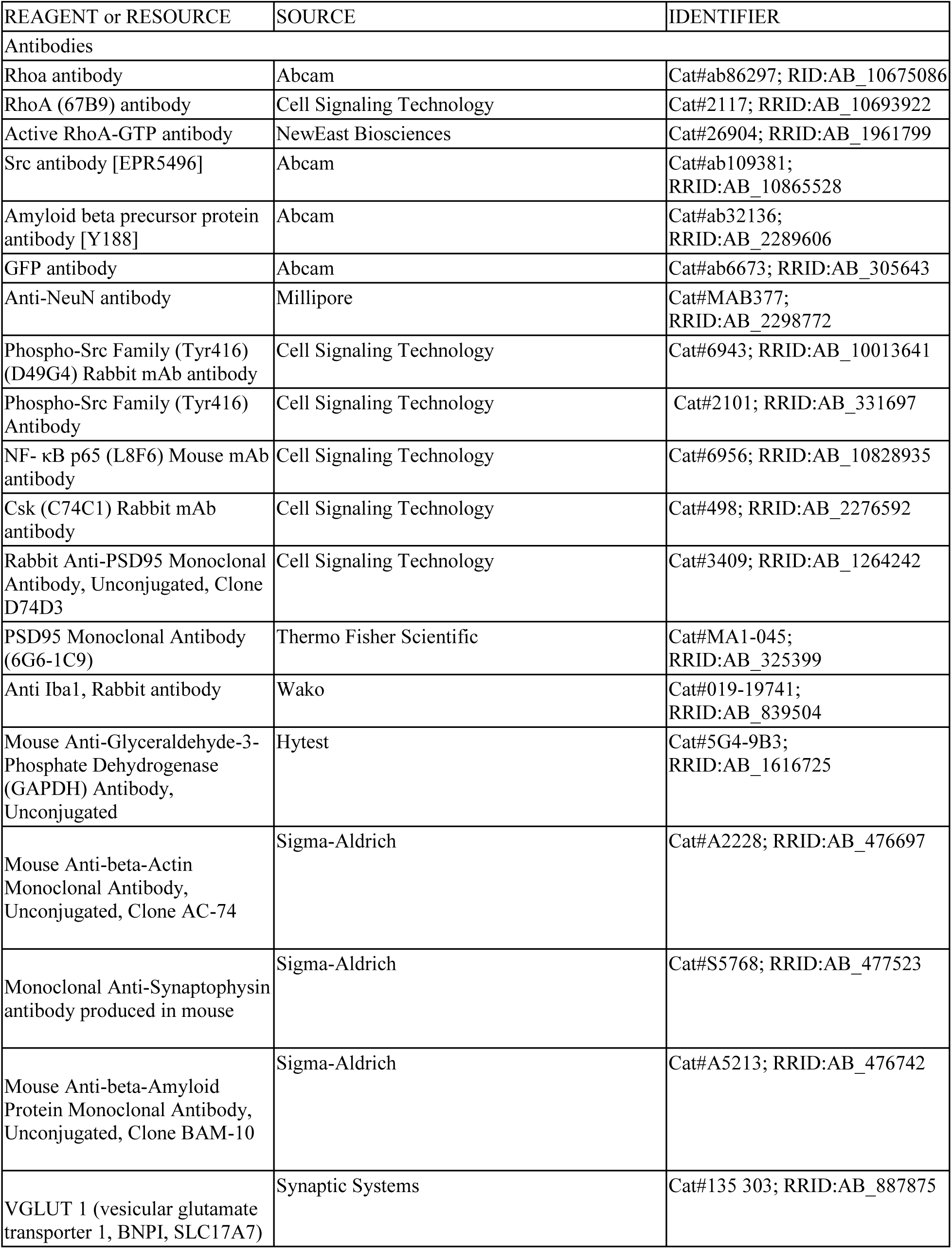
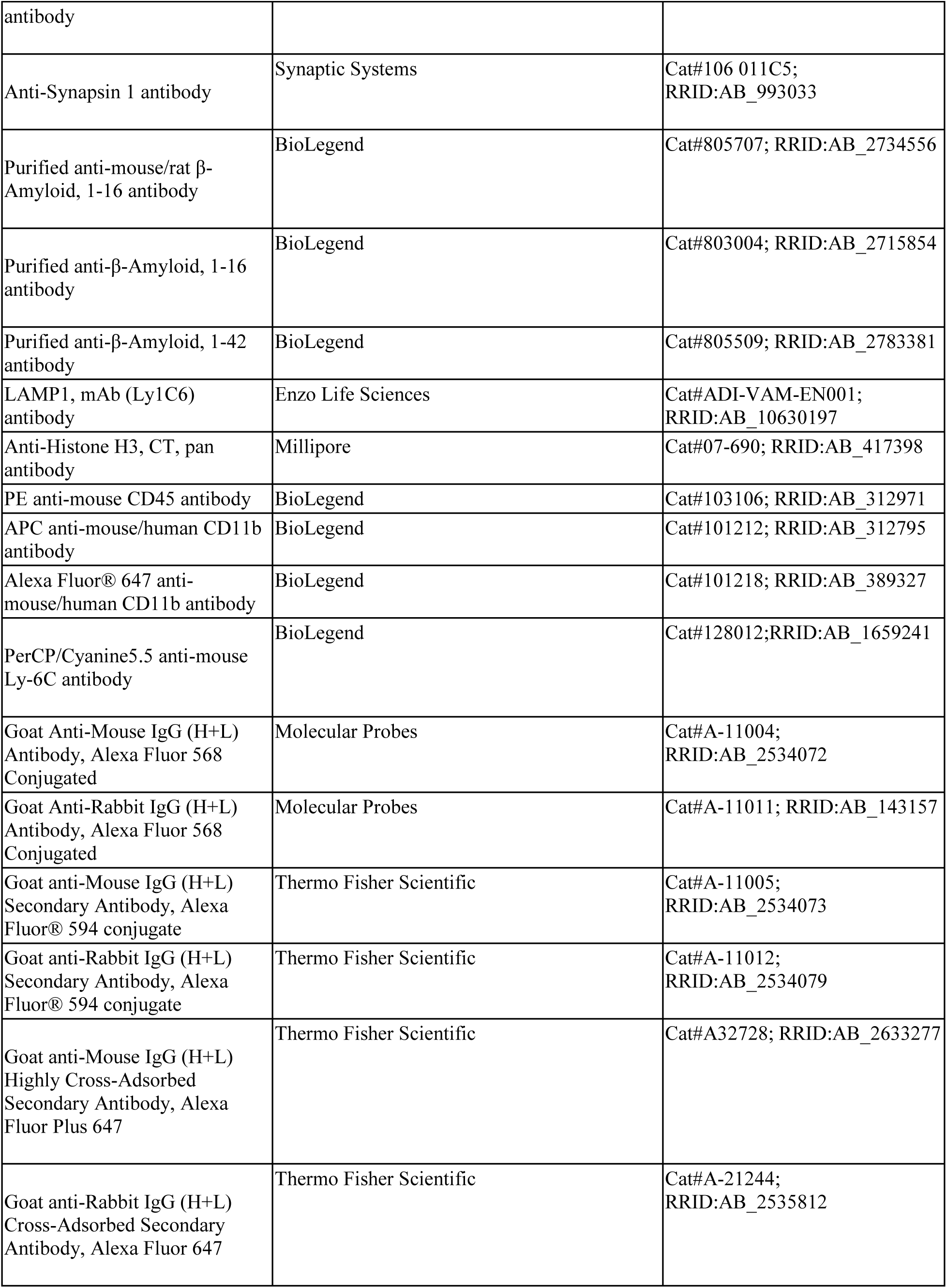
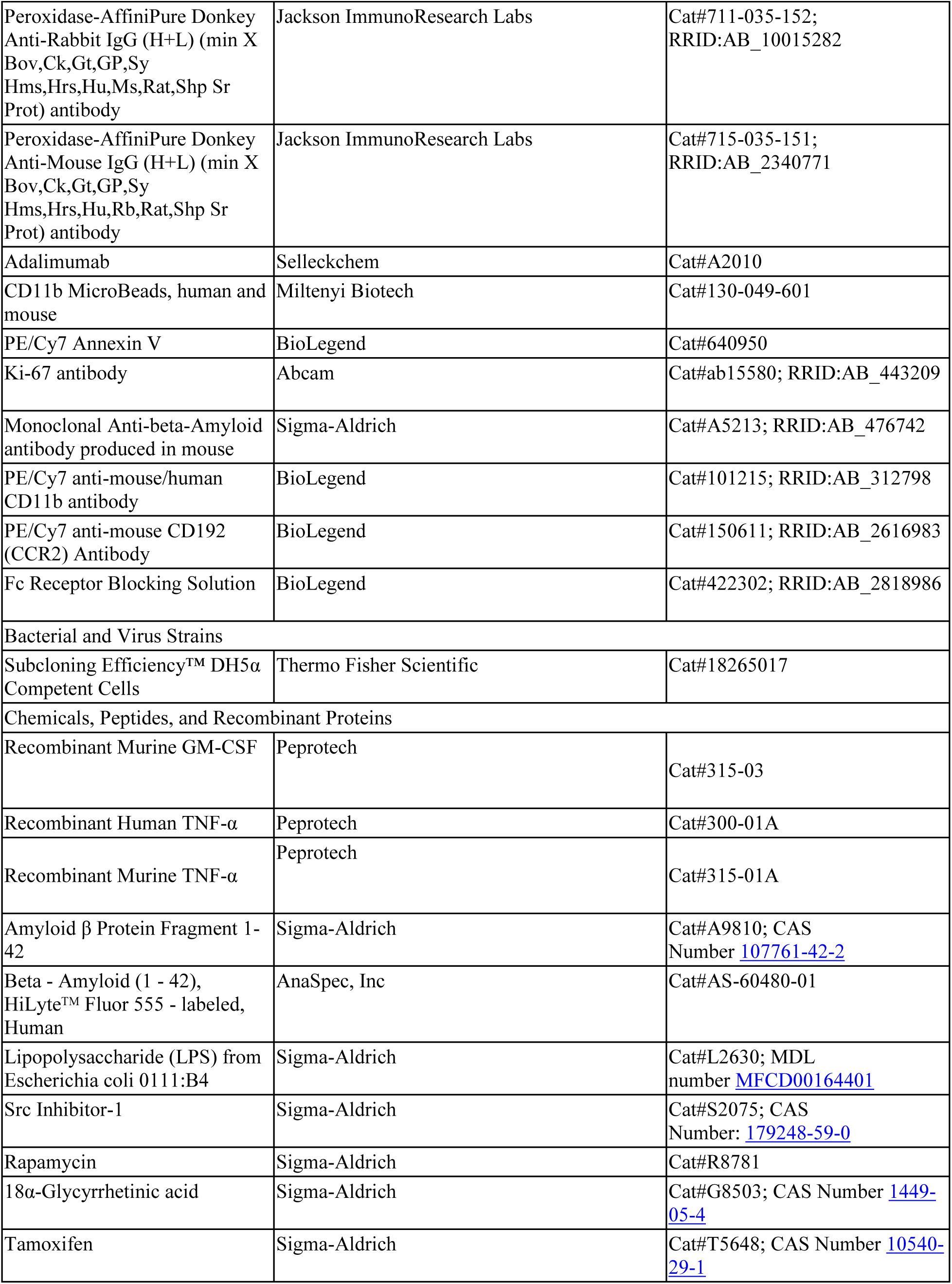
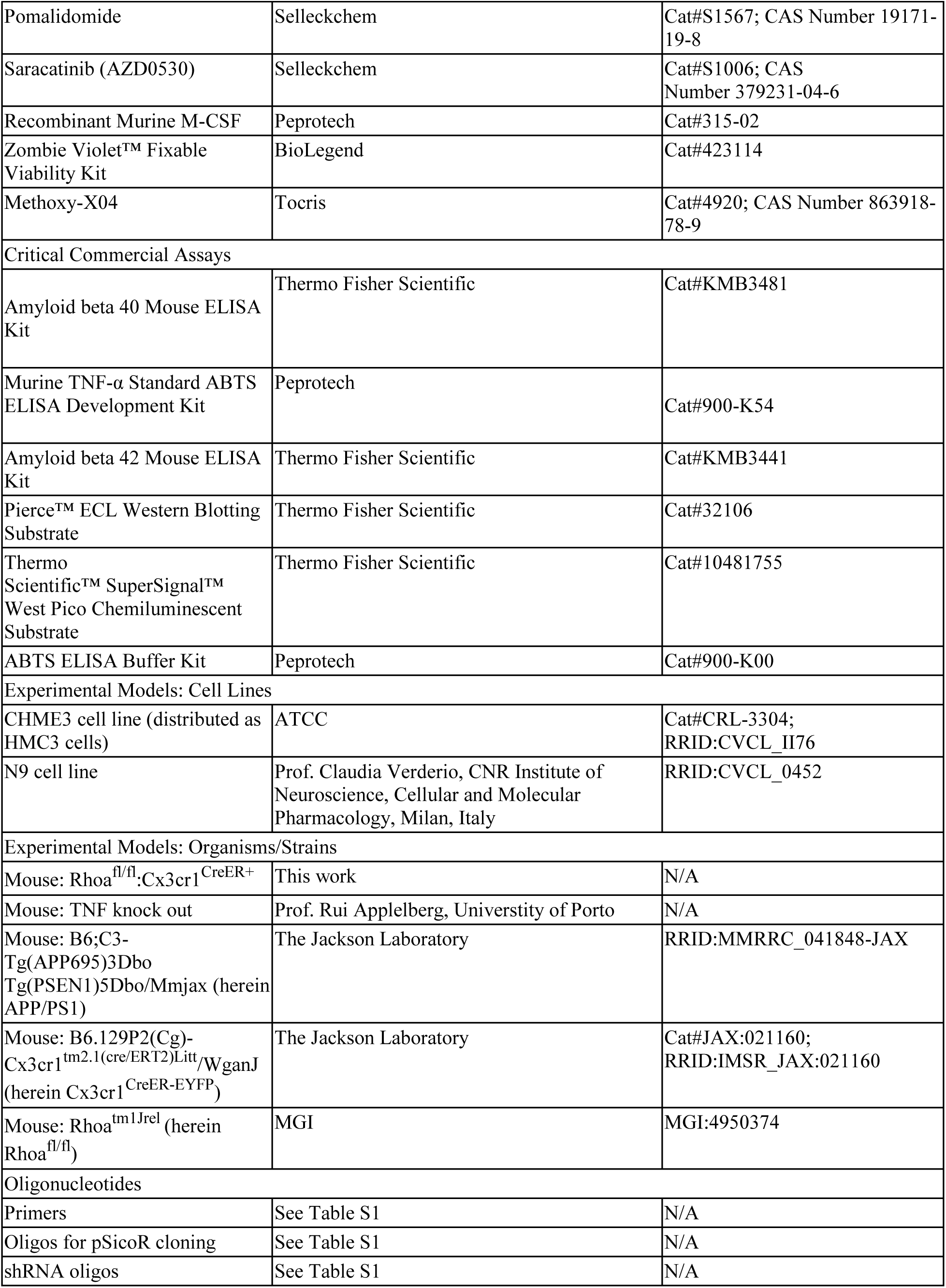
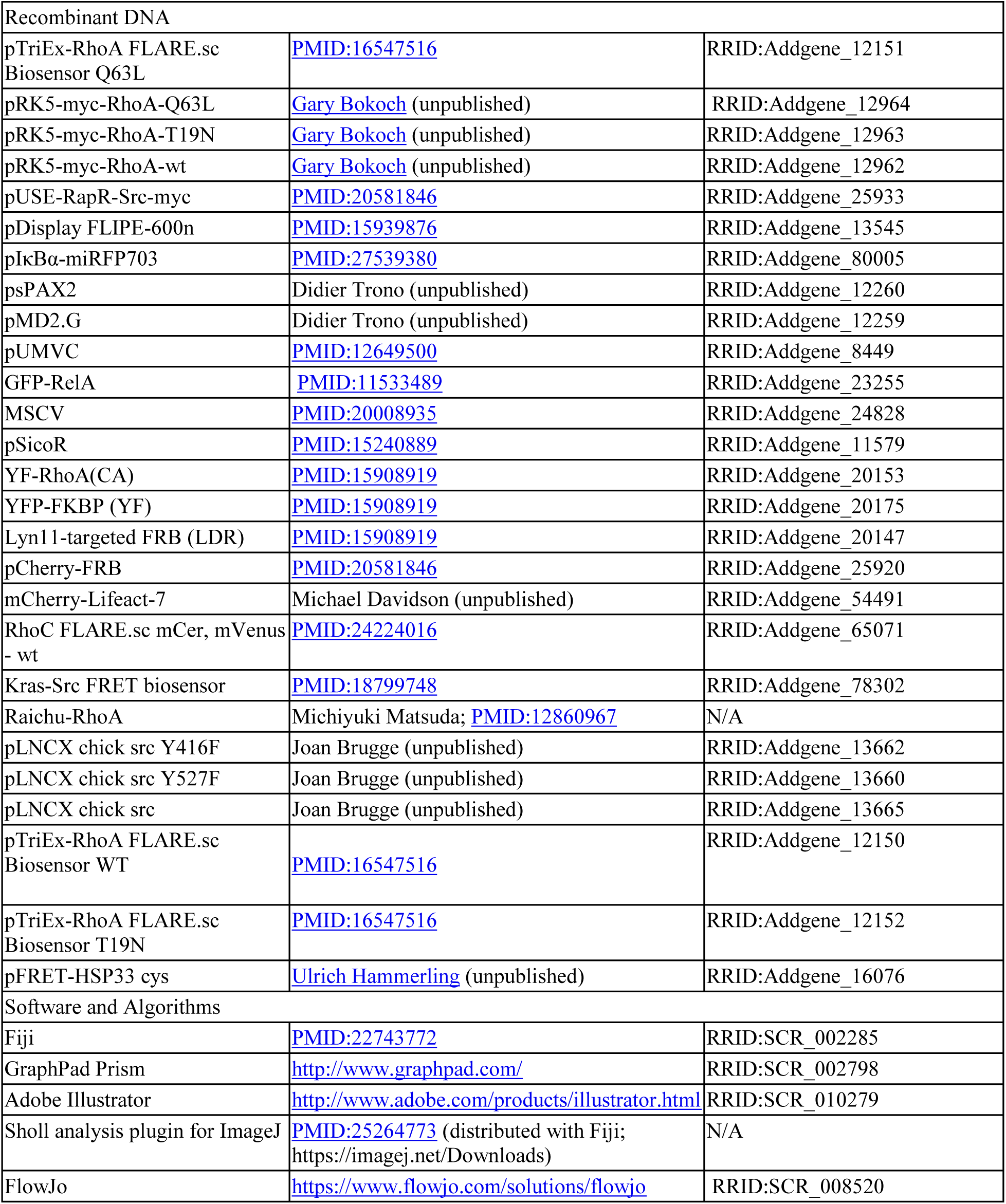

## KEY RESOURCES TABLE

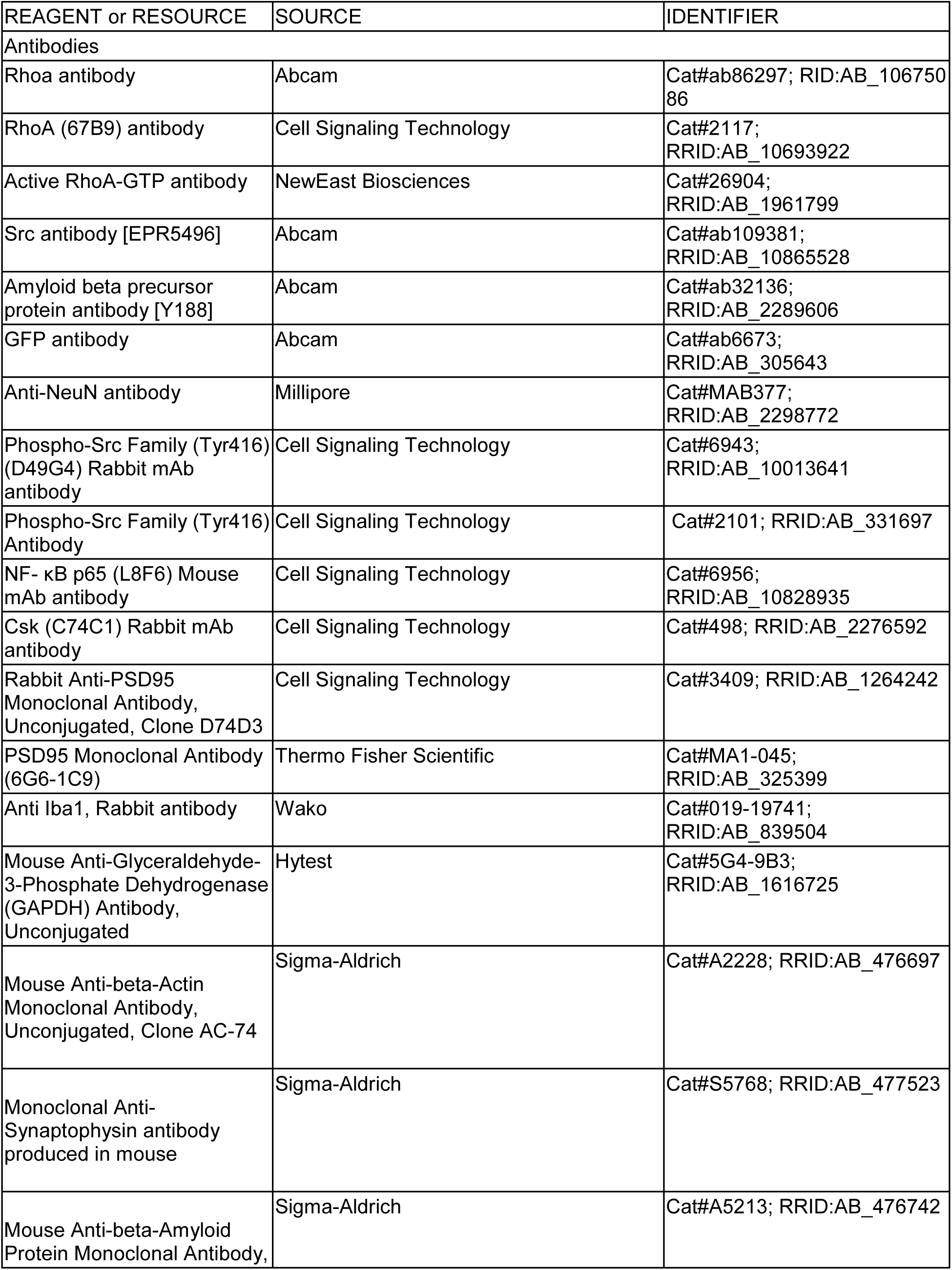
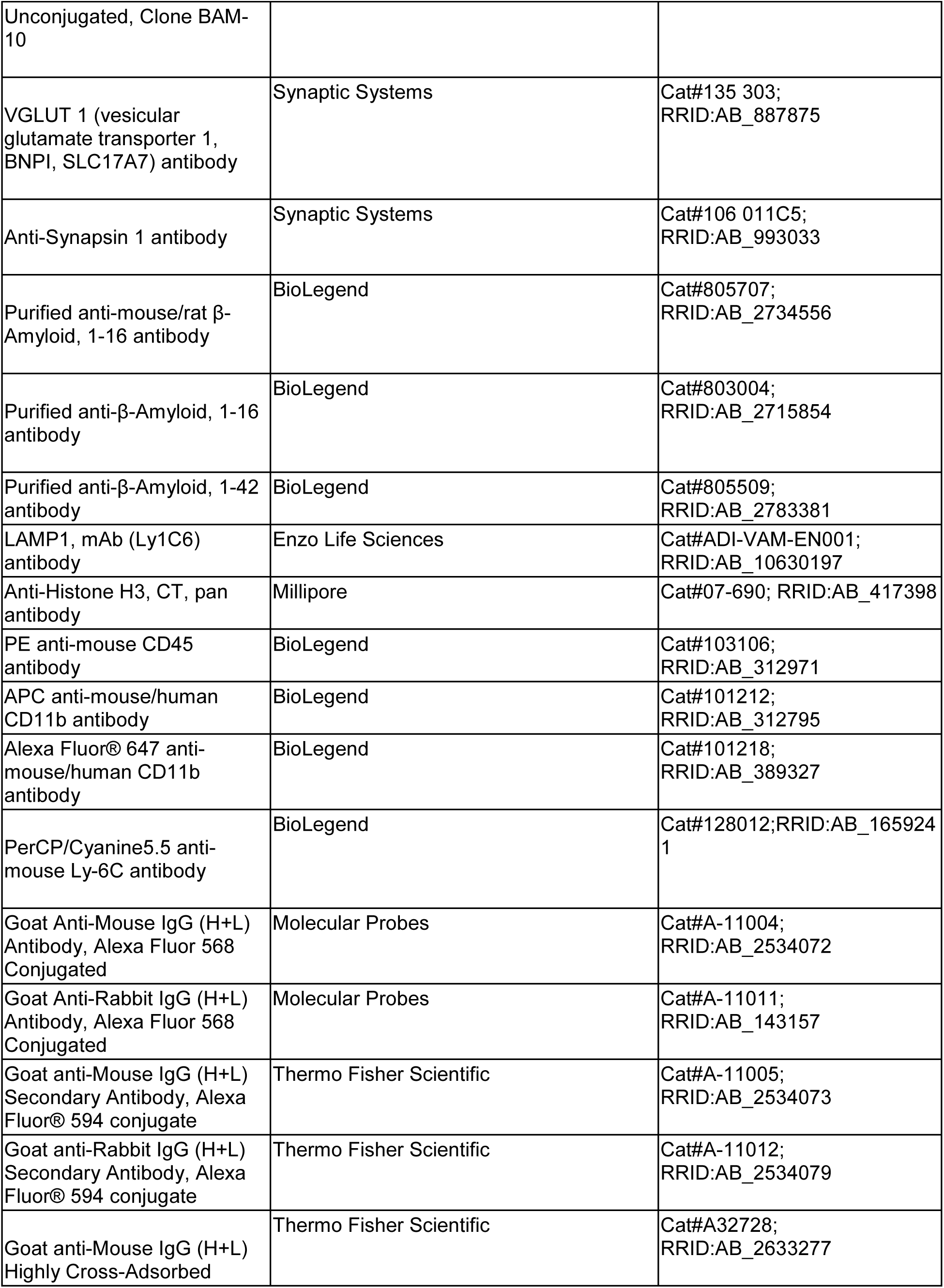
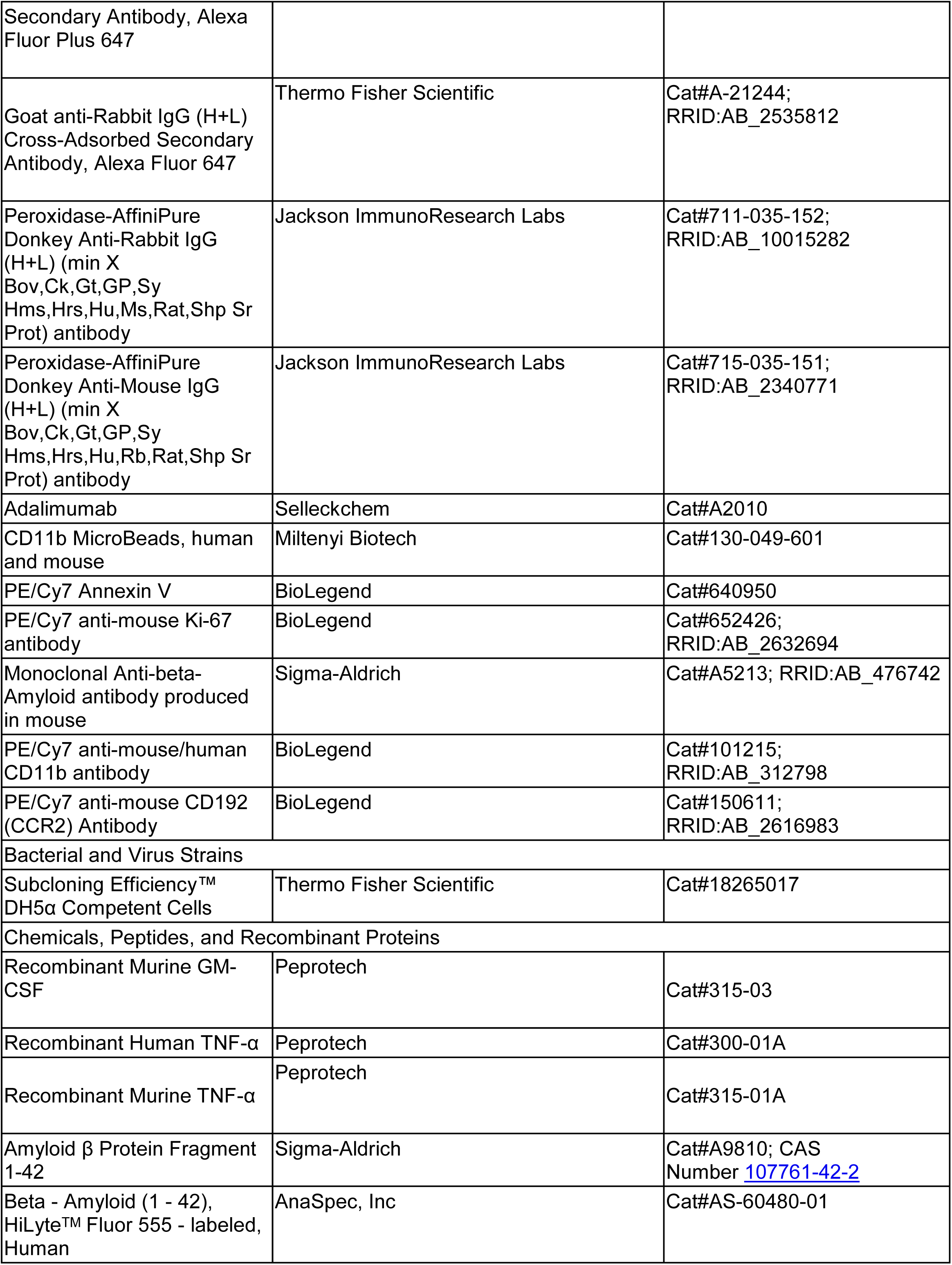
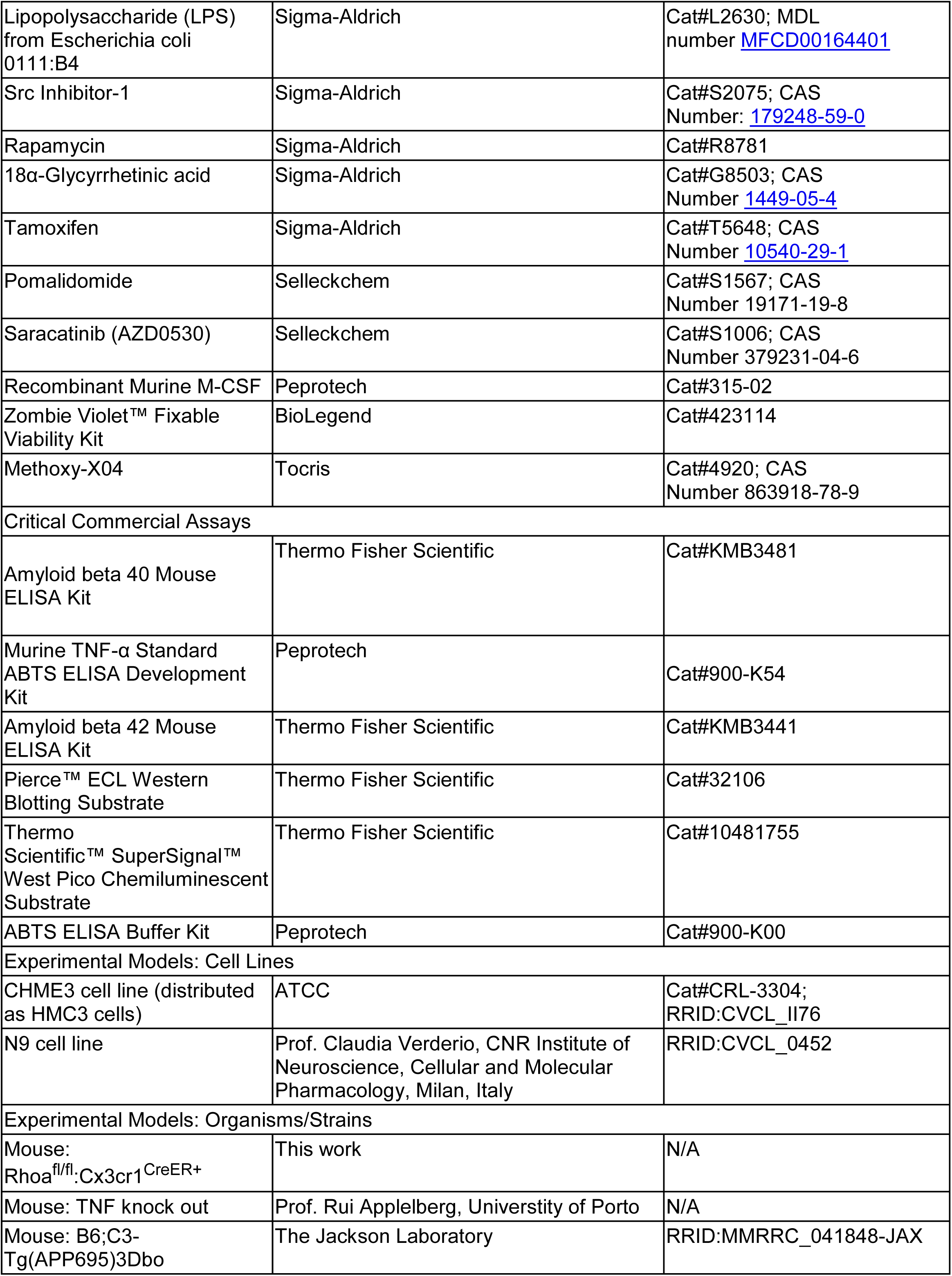
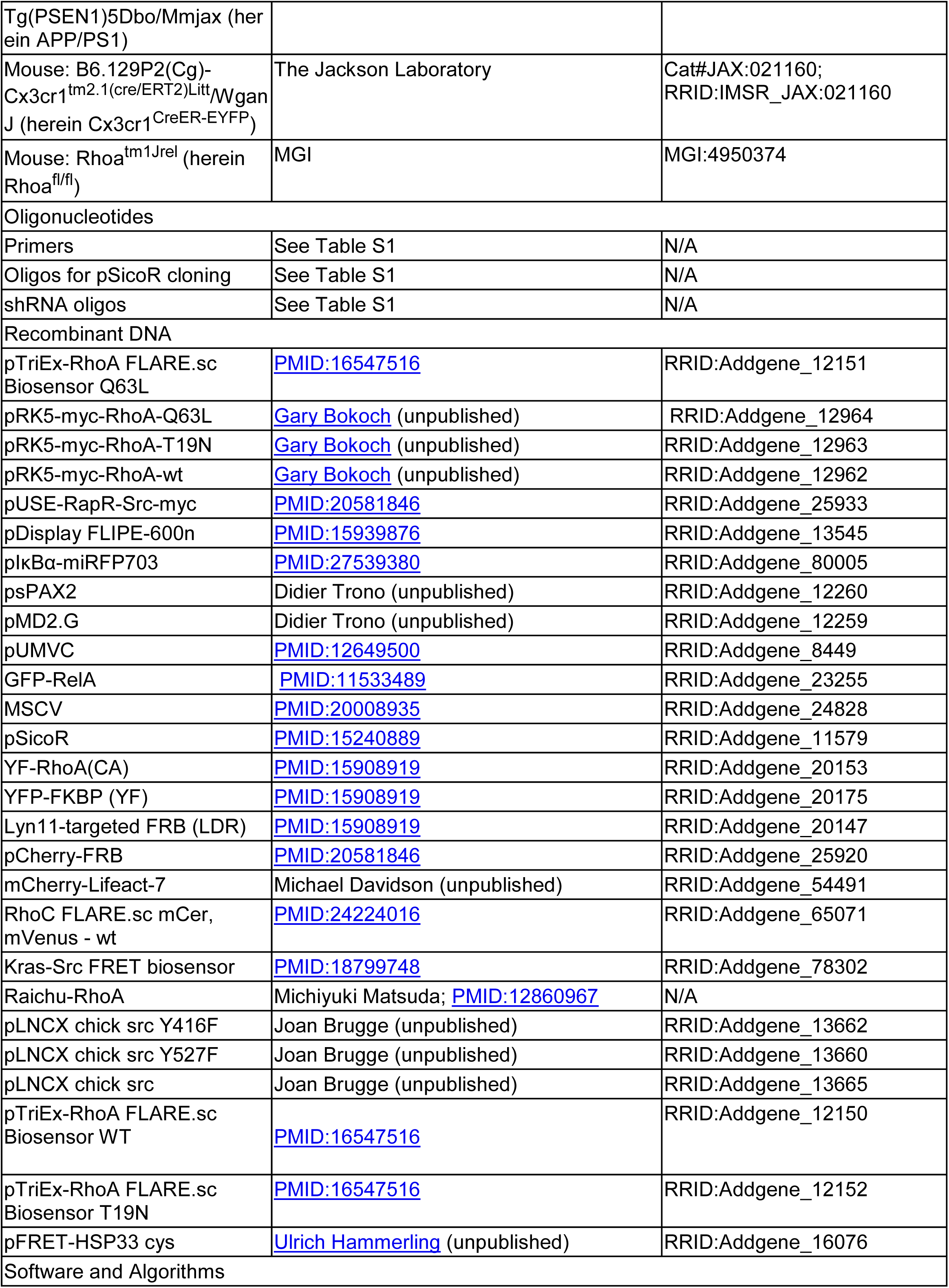
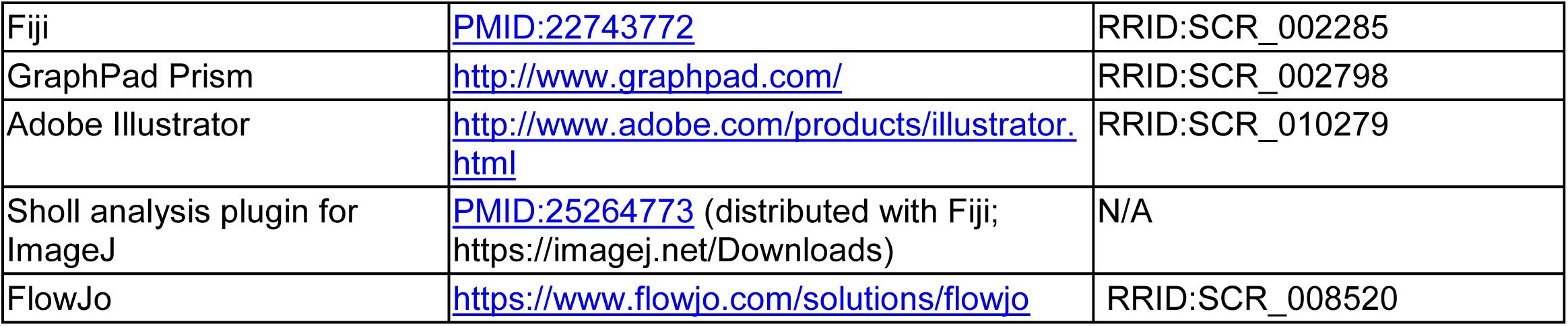

## Supplemental Text and Figures

**Supplemental Figure 1.**
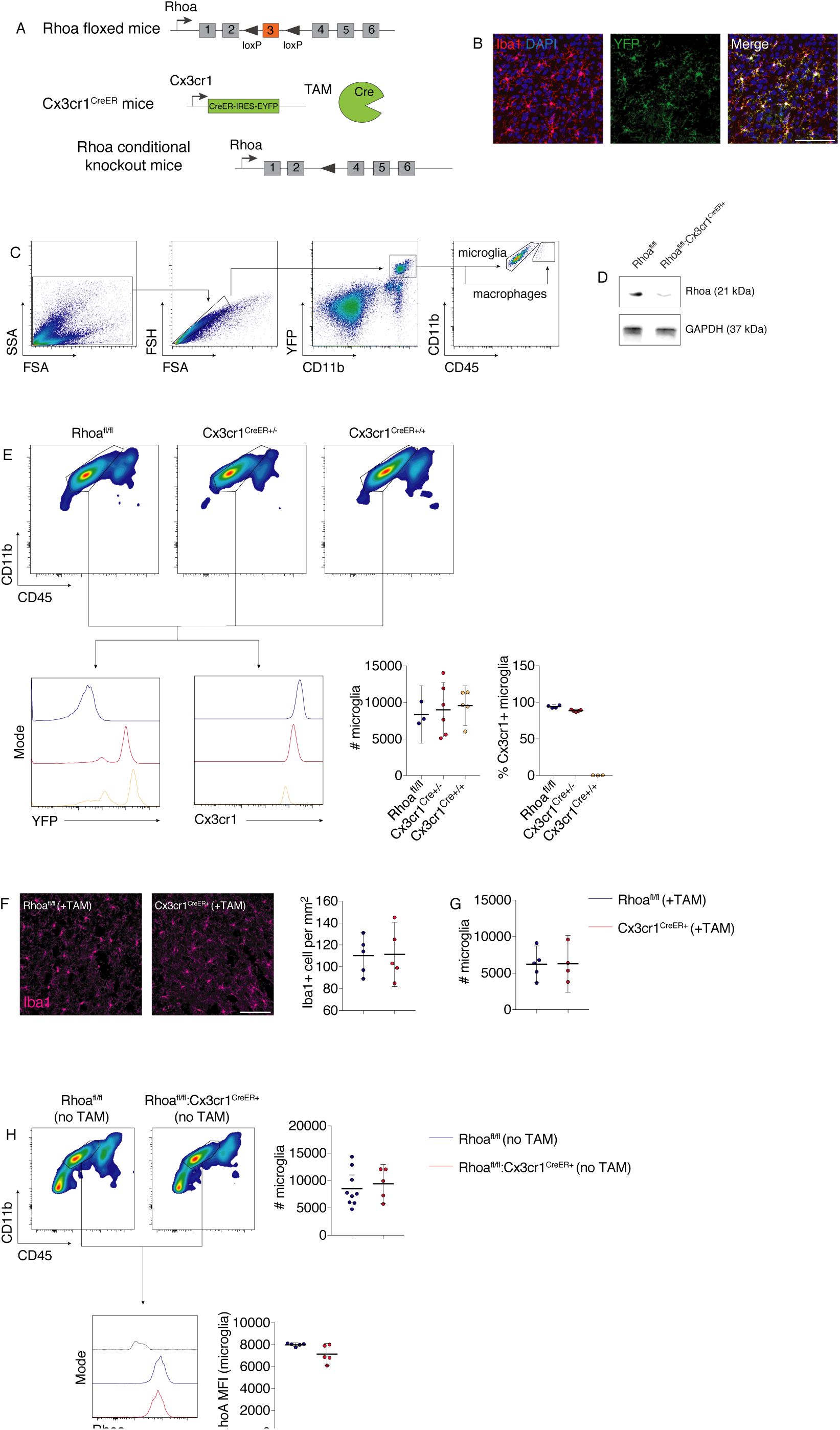
Ablation of Rhoa in microglia and Cx3cr1 haploinsufficiency does not impact microglial Rhoa expression and microglial numbers (Related to Fig. 1). **A**, breeding scheme for tamoxifen-inducible microglia-specific *Rhoa* gene inactivation in mice. Crossing mice bearing CreER-IRES-EYFP transgene within the Cx3cr1 locus (top) with mice in which the exon 3 of the *Rhoa* gene is flanked by loxP sites (middle) gives rise to mice in which Rhoa deletion in microglia/macrophage is achieved by tamoxifen administration (bottom). **B**, confocal imaging of brain sections from Cx3cr1-EYFP-Cre^ER+^ heterozygous mice immunolabeled using anti-Iba1 antibody (red) and anti-YFP antibody (green). Virtually all YFP^+^ cells were also Iba1^+^ (images are representative of 4 mice). Scale bar: 100 μm. **C**, Gating strategy for sorting microglia from the brains of Cx3cr1-EYFP-Cre^ER+^ mice. More than 95% of the EYFP^high^ gated population in the brain corresponded to microglia (CD45^mid^CD11b^+^), the remaining cells were composed of non-parenchymal brain macrophages (CD45^high^CD11b^+^). **D**, Western blot for Rhoa on lysates from MACS-separated microglia collected from the brains of Rhoa^fl/fl^ and Rhoa^fl/fl^:Cx3cr1^CreER+^ mice 90 days after TAM administration. GAPDH was the loading control (images are representative of 3 mice per condition). **E**, Flow cytometry analysis of Cx3cr1 on microglia from Rhoa^fl/fl^, Cx3cr1^CreER+/-^ and Cx3cr1^CreER+/+^ mice (n=5 mice per condition). Mice from these experiments were not given TAM. Dot plots are mean and 95% confidence interval and significance was analyzed by One-way ANOVA. Cell debris was excluded by size. **F and G**, confocal imaging of Iba1 immunostaining on brain sections (F) and flow cytometry analysis of microglial numbers (G) in Rhoa^fl/fl^ and Cx3cr1^CreER+^ mice 21 days after TAM (n=5 mice per condition). Dot plots are mean and 95% confidence interval and significance was analyzed by Mann-Whitney test. Scale bar: 100 μm. **H**, Flow cytometry analysis of Rhoa on microglia from Rhoa^fl/fl^ and Rhoa^fl/fl^:Cx3cr1^CreER+^ mice (n=7 mice per condition). Mice from these experiments were not given TAM. Dot plots are mean and 95% confidence interval and significance was analyzed by Mann-Whitney test. Cell debris was excluded by size.

**Supplemental Figure 2.**
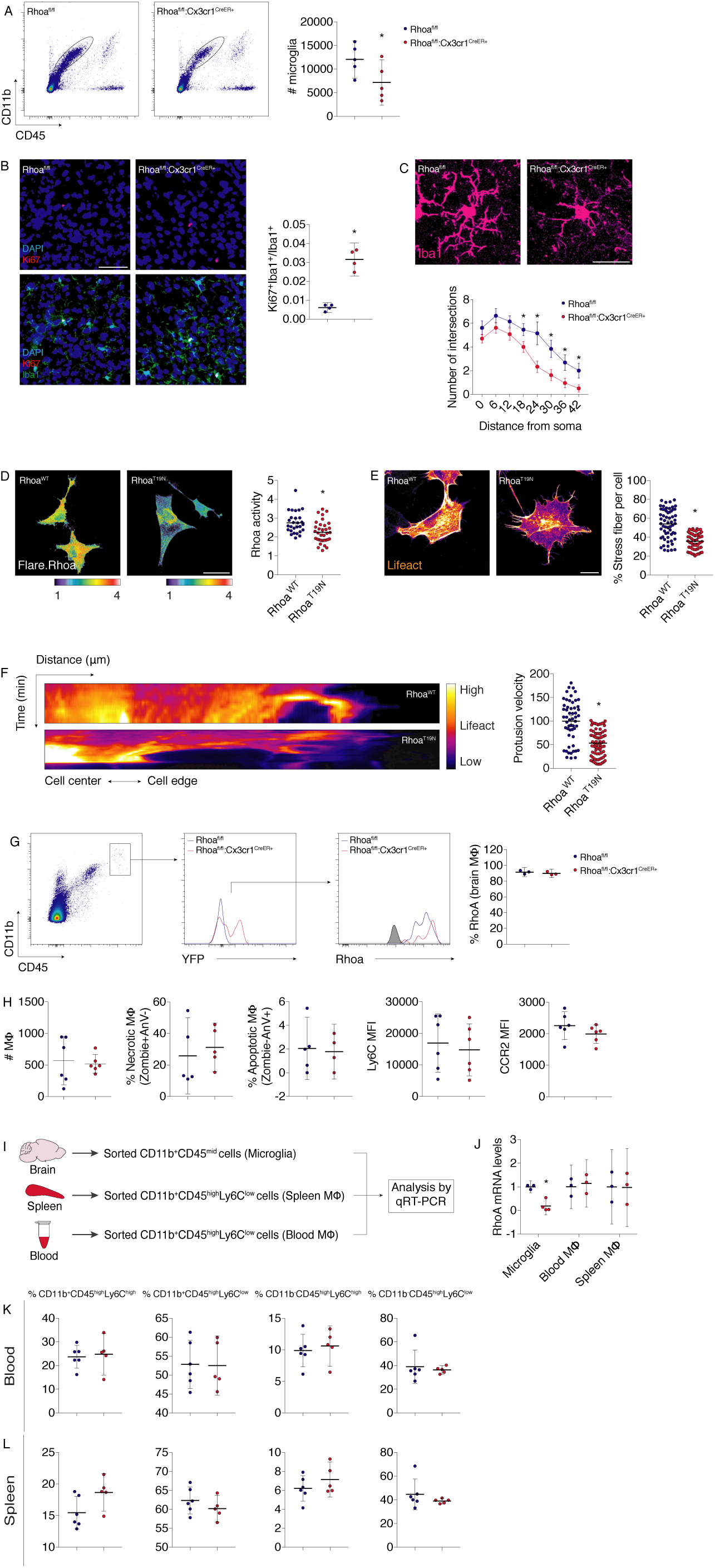
Microglia morphology changes correlates with alterations in actin cytoskeleton organization and dynamics and macrophage populations in Rhoa^fl/fl^:Cx3cr1^CreER+^ mice (Related to Fig. 1). **A**, Flow cytometry analysis of microglial numbers in Rhoa^fl/fl^ and Rhoa^fl/fl^:Cx3cr1^CreER+^ mice 150 days after TAM administration (n=5 mice per condition). Dot plots show microglia cell counts (mean and 95% confidence interval). Cell debris was excluded by size. *P<0.05 (Mann-Whitney test). **B,** histological confocal analysis for Iba1 and Ki67 on tissue sections from the neocortex of Rhoa^fl/fl^ and Rhoa^fl/fl^:Cx3cr1^CreER+^ mice 150 days after TAM administration (n=4 mice per condition). Dot plots are mean and 95% confidence interval). *P<0.05 (Mann-Whitney test). Scale bar: 50 μm. **C,** histological confocal analysis for Iba1 on tissue sections from the neocortex of Rhoa^fl/fl^ and Rhoa^fl/fl^:Cx3cr1^CreER+^ mice 150 days after TAM administration (n=5 mice for each time point). Dot plots (mean and 95% confidence interval) depicting Sholl analysis for each condition is shown. *P<0.05 (Two-way ANOVA comparing Rhoa^fl/fl^ vs. Rhoa^fl/fl^:Cx3cr1^CreER+^ for each distance). Scale bar: 20 μm. **D-F**, Representative images (D and E) or kymograms (F) of CHME3 microglial cultures expressing a Rhoa FRET biosensor (Flare.Rhoa, D) or Lifeact (E and F) transfected with wild-type (WT) Rhoa or dominant-negative (T19N) Rhoa mutant (n=25 cells per condition from 2 independent experiments in D, n=60 cells per condition from 2 independent experiments in E, n=54-80 protrusions polled across 14-20 cells from 2 independent experiments F). Dot plots (means and 95% confidence interval) display FRET/CFP ratio images (D), stress fibers content (E) and microglial process velocity (F). Pseudocolor ramps represent min/max FRET/CFP ratios (D) and lifeact intensity (E and F). *P<0.05 (unpaired t test) vs. Rhoa^WT^. Scale bars: 20 μm (D), 10 μm (E). **G and H**, Flow cytometry analysis of Rhoa, cell death and reactivity markers in brain macrophages from Rhoa^fl/fl^ and Rhoa^fl/fl^:Cx3cr1^CreER+^ mice 150 days after TAM administration (n=3-6 mice). Dot plots are mean and 95% confidence interval. Significance was analyzed by Mann-Whitney test. Cell debris was excluded by size. **I and J**, mRNA was harvested from flow cytometry-sorted brain microglia, blood monocytes or spleen macrophages from Rhoa^fl/fl^ and Rhoa^fl/fl^:Cx3cr1^CreER+^ mice 35-45 days after TAM administration. qRT-PCR determined the abundance of *Rhoa* mRNA transcripts in each population (n=3-4 mice). Dot plots displays means and 95% confidence interval. *P<0.05 (Mann-Whitney test). **K and L**, FACS analysis showing the percentage of different CD45^high^ populations in the blood and spleen of Rhoa^fl/fl^ and Rhoa^fl/fl^:Cx3cr1^CreER+^ mice 35-45 days after TAM administration (n=5-6 mice). Dot plots depict percent cell counts (mean and 95% confidence interval) for each cell population. Significance was analyzed by Mann-Whitney test.

**Supplemental Figure 3.**
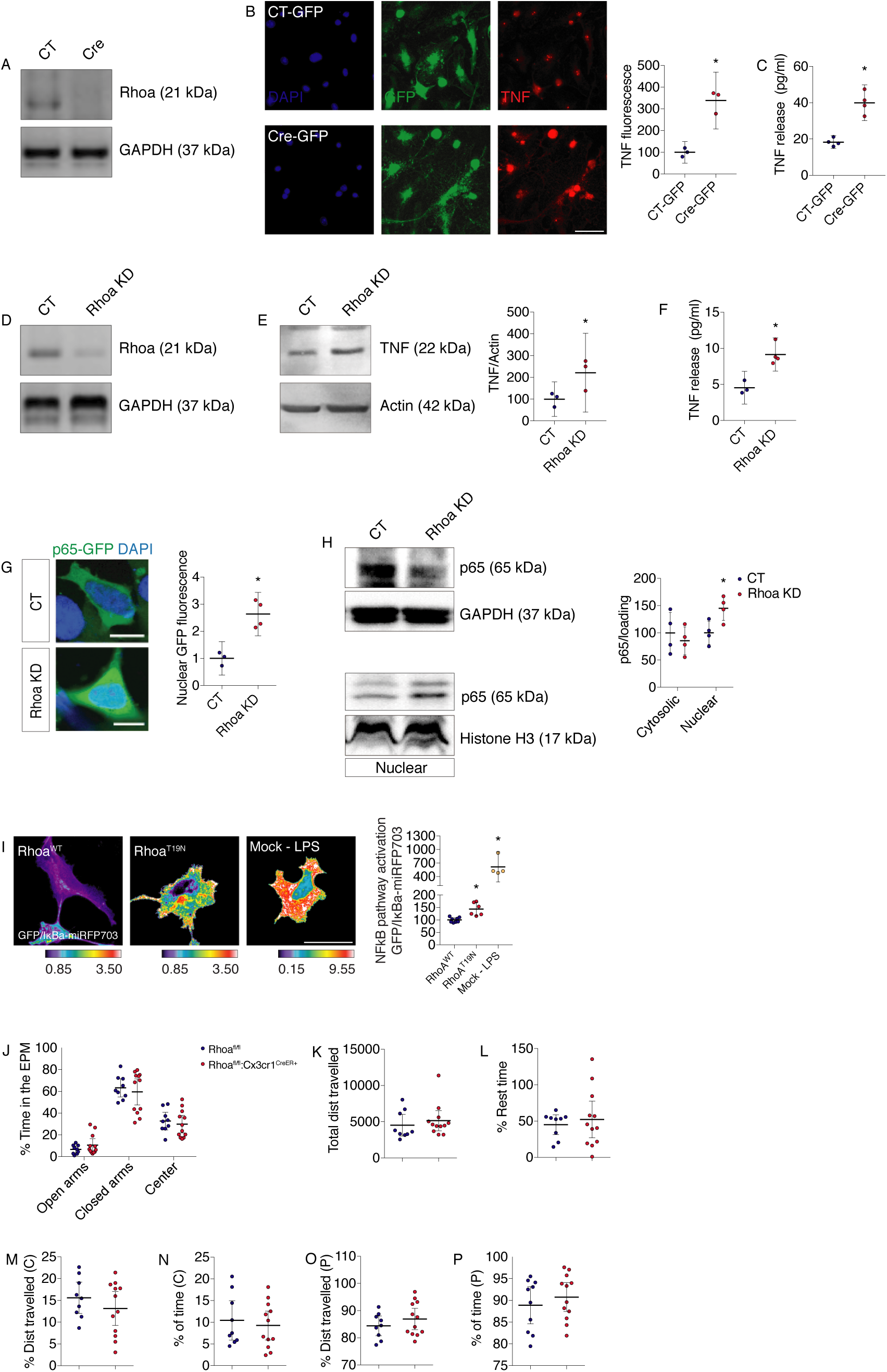
Rhoa depletion leads to cell autonomous microglial activation and behavior phenotype of Rhoa^fl/fl^:Cx3cr1^CreER+^ mice. (Related to Fig. 1 and 2). **A-C**, Western blot for Rhoa (A), immunocytochemistry for Tnf (B) and ELISA on the culture media of primary cortical microglial cell cultures from Rhoa floxed mice (n=3-4 independent cultures). Cultures were transduced with lentiviruses coding GFP (CT-GFP) or Cre recombinase (Cre-GFP). Dot plots (means and 95% confidence interval) displays Rhoa expression, Tnf fluorescence or Tnf content in pg/ml. *P<0.05 (unpaired t test). Scale bar: 20 μm. **D and E**, Western blot for Rhoa and Tnf on cell lysates of Rhoa shRNA stable (Rhoa KD) or control (CT; pLKO) CHME3 microglial cell sub-clones (n=3 independent cultures). GAPDH and actin were used as the loading control. Dot plots (means and 95% confidence interval) displays Tnf per actin ratio. *P<0.05 (unpaired t test). **F**, ELISA on the culture media of Rhoa shRNA stable (Rhoa KD) or control (CT; pLKO) N9 microglial cell sub-clones (n=3-4 independent cultures). Dot plots (means and 95% confidence interval) displays Tnf content in pg/ml. *P<0.05 (Mann-Whitney test). **G**, Confocal imaging of Rhoa shRNA stable (Rhoa KD) or control (CT; pLKO) N9 microglial cell sub-clones transfected with GFP-tagged p65 subunit of the NFκB complex and stained with DAPI (n=3-4 independent cultures). Dot plots (means and 95% confidence interval) displays nuclear GFP fluorescence normalized to the CT values. *P<0.05 (Mann-Whitney test). Scale bars, 10 μm. **H**, Western blots for p65 subunit of the NFκB complex on cytosolic and nuclear extracts of Rhoa shRNA stable (Rhoa KD) or control (CT; pLKO) N9 microglial cell sub-clones (n=4 independent cultures). GAPDH and histone H3 were used as loading controls. *P<0.05 (Mann-Whitney test). **I**, Representative images of CHME3 microglial cultures expressing an NFkB pathway activity biosensor (IkBa-miRFP703) transfected with GFP-Rhoa WT or GFP-Rhoa T19N (n= 4-8 cells from 2 independent experiments). LPS (1 µg/ml; 24 h) was used as positive control for IkBa degradation. Dot plots (means and 95% confidence interval) displays GFP/miRFP703 ratio images coded according to the pseudocolor ramp. *P<0.05 (One-way ANOVA) vs. Rhoa WT. Scale bar: 10 μm. **J-P**, Rhoa^fl/fl^ and Rhoa^fl/fl^:Cx3cr1^CreER+^ mice were evaluated in the elevated-plus maze (EPM, J) and in the open field arena (K-P) 35-45 days after TAM administration (n=9-12 mice). No alteration in general locomotor activity or anxiety-related behavior (no significant difference in the time spent in the open arms (OA), in the closed arms (CA) or in the center of the maze) was observed between genotypes. Dot plots are mean and 95% confidence interval; significance was analyzed by unpaired t test.

**Supplemental Figure 4.**
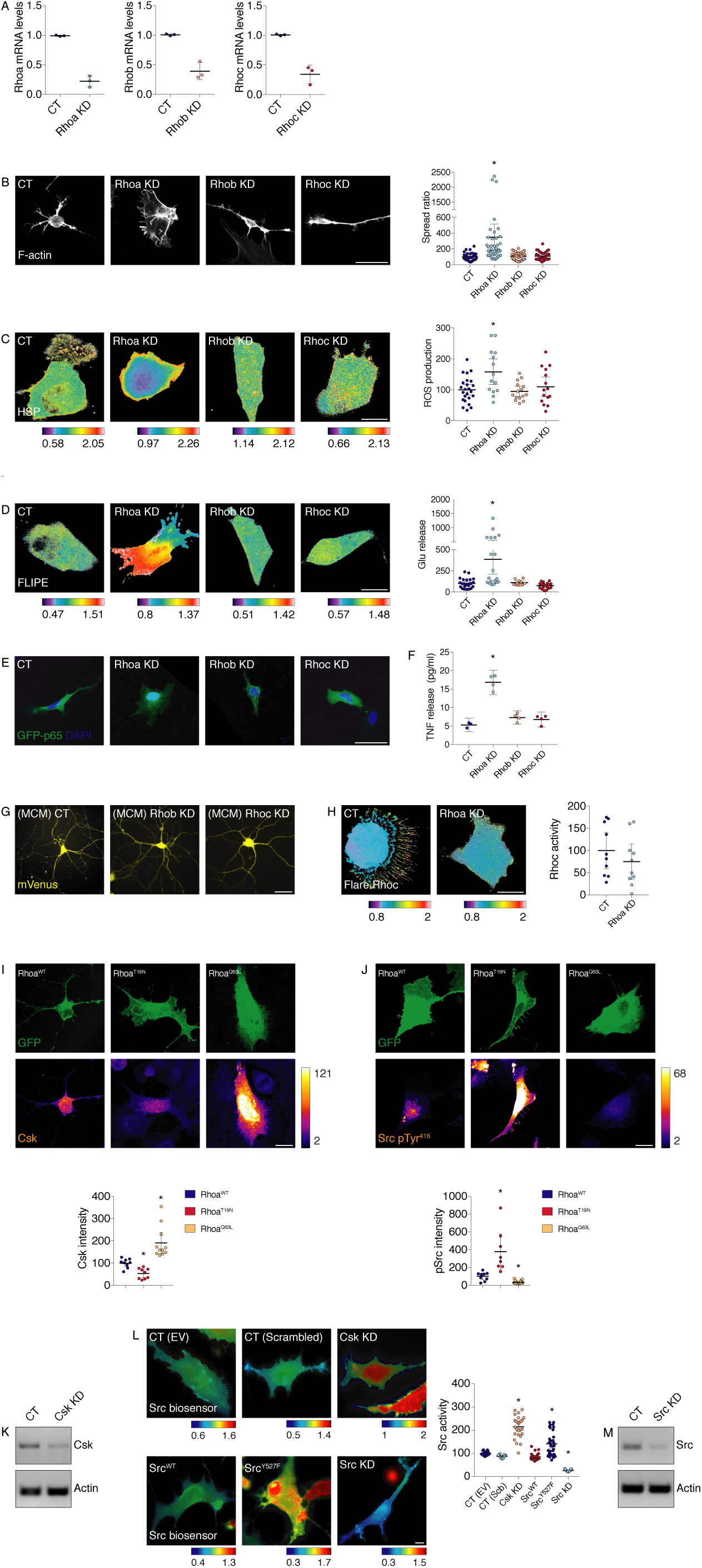
Knockdown of Rhob or Rhoc does not trigger spontaneous microglia activation and Rhoa regulation of Csk/Src pathway (Related to Fig. 2 and 3). **A**, qRT-PCR for Rhoa, Rhob and Rhoc in RNA samples harvested from primary microglial cultures (n=3 independent cultures). Cultures were transduced with lentiviruses coding pLKO (CT), Rhoa shRNA, Rhob shRNA or Rhoc shRNA. Dot plots are mean and 95% confidence interval. **B**, fluorescence imaging of Lifeact (labeling f-actin) in CHME3 microglial cell cultures previously infected with lentiviruses coding pLKO (CT), Rhoa shRNA, Rhob shRNA or Rhoc shRNA (n=39-44 cells from 3 independent cultures). Dot plots (means and 95% confidence interval) displays the microglia spread ratio (a measure of cell morphology) normalized to the CT. *P<0.05 (Kruskal-Wallis test) vs. CT. Scale bar: 20 μm. **C and D**, primary microglial cultures expressing the ROS (HSP; C) or the glutamate release (FLIPE; D) FRET biosensor, were infected with lentiviruses coding pLKO (CT), Rhoa shRNA, Rhob shRNA or Rhoc shRNA (n=14-24 (C) and 11-24 (D) cells from 3 independent experiments). Time-lapse CFP/FRET ratio images were averaged for 10 min and plotted. Pseudocolor ramps represent Donor/FRET min/max ratios. Dot plots (mean and 95% confidence interval) were normalized to CT. *P<0.05 (One-way ANOVA) vs. CT. Scale bars: 5 μm. **E**, confocal imaging of GFP-tagged p65 subunit of the NFκB complex (co-stained with DAPI) in CHME3 microglial cell cultures previously infected with lentiviruses coding pLKO (CT), Rhoa shRNA, Rhob shRNA or Rhoc shRNA (n=5 independent cultures per condition). Scale bar: 20 μm. **F**, ELISA on the culture media of shRNA (CT), Rhoa shRNA, Rhob shRNA or Rhoc shRNA N9 microglial cell sub-clones (n=3-4 independent cultures). Dot plots (means and 95% confidence interval) displays Tnf content in pg/ml. *P<0.05 (One-way ANOVA). **G**, primary hippocampal neuronal cultures expressing the yellow fluorescent protein variant mVenus were incubated for 24 h with conditioned media from primary microglial cultures (MCM) that were previously infected with pLKO (CT), Rhob shRNA or Rhoc shRNA lentiviruses (n=3 independent hippocampal cultures). Scale bar: 50 μm. **H**, primary cortical microglial cultures expressing the Flare.RhoC FRET biosensor were infected with lentiviruses coding pLKO (CT) or Rhoa shRNA (n=10 cells for each condition from 3 independent experiments). Time-lapse FRET/CFP ratio images were averaged for 10 min and plotted. Pseudocolor ramps represent FRET/donor min/max ratios. Dot plots (mean and 95% confidence interval) were normalized to CT. *P<0.05 (unpaired t test). Scale bar: 5 μm. **I and J**, Confocal imaging of CHME3 microglia transfected with GFP-tagged WT Rhoa, T19N Rhoa or Q63L Rhoa and immunolabeled for Csk of Src pTyr416 (n=4 independent cultures). Dot plots (means and 95% confidence interval) displays Csk or pSrc fluorescence normalized to the Rhoa^WT^ values. *P<0.05 (One-way ANOVA). Scale bars, 10 μm. **K**, Western blot for Csk on cell lysates of Csk shRNA stable (Csk KD) or control (CT; pLKO) CHME3 microglial cell sub-clones (n=3 independent cultures). Actin was used as the loading control. **L**, Representative images of CHME3 microglial cultures expressing a FRET-based Src activity biosensor (KRas Src YPet) transfected with empty vector (EV), scrambled vector, Csk shRNA (Csk KD), WT Src, constitutively active (Y527F) Src mutant or Src shRNA (Src KD) (n is from at least 13 cells per condition from 2 independent experiments). Dot plots (means and 95% confidence interval) display CFP/FRET ratio images normalized to EV. Pseudocolor ramps represent min/max CFP/FRET ratios. *P<0.05 (Kruskal-Wallis test) vs. CT (EV). Scale bars: 20 μm. **M**, Western blot for Src on cell lysates of Src shRNA stable (Src KD) or control (CT; pLKO) CHME3 microglial cell sub-clones (n=3 independent cultures). Actin was used as the loading control.

**Supplemental Figure 5.**
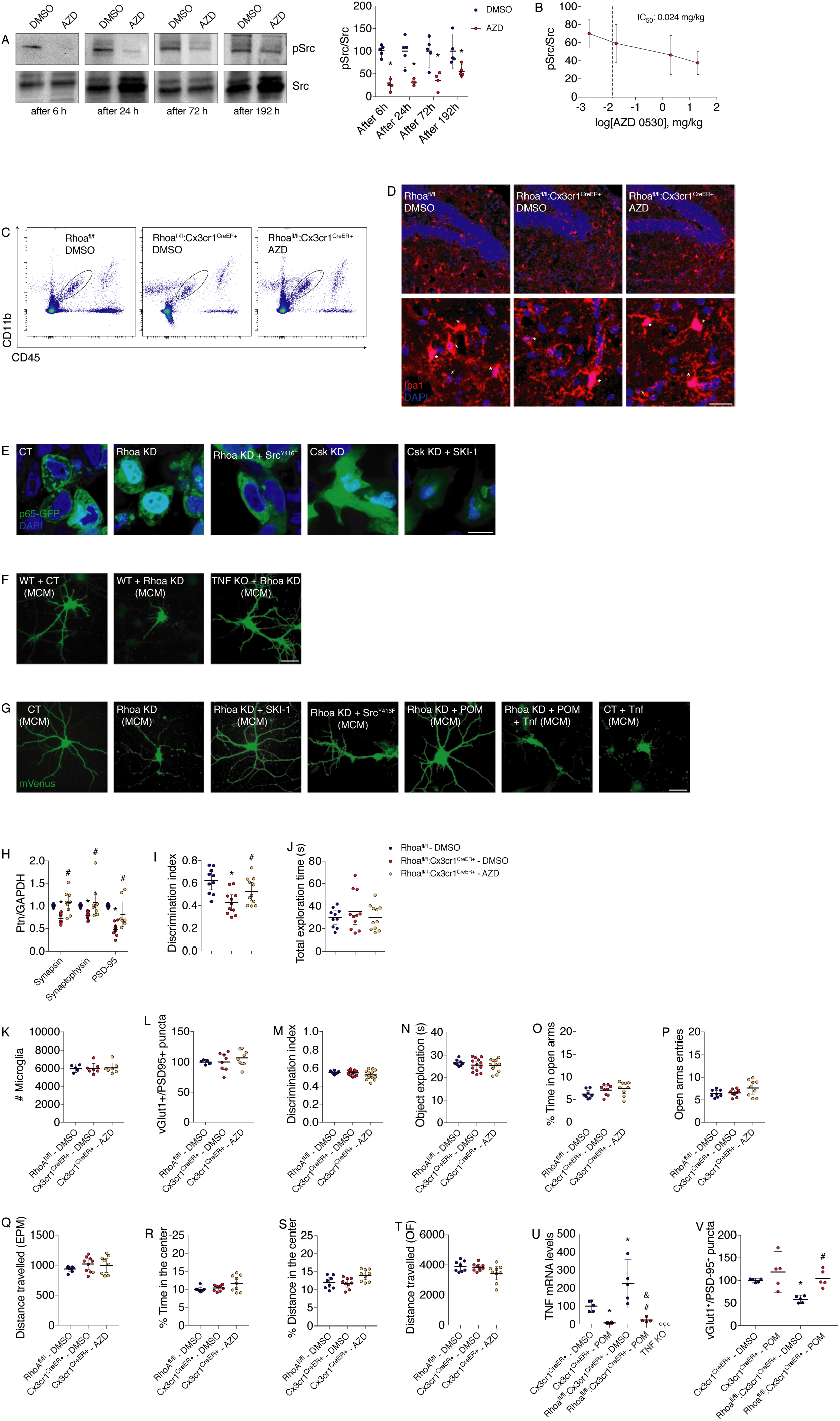
Rhoa regulation of Src-to-Tnf signaling in microglia impacts neurotoxicity (Related to Fig. 4). **A**, Western blot for Src pTyr^416^ on lysates from the brains of naïve WT mice injected IP with a single dose of DMSO (n=5 mice per condition) or 10 mg/kg AZD 0530 (n=4-5 mice per condition). Brains were harvested 6 h, 24 h, 72 h or 196 h after injections. Src was used as the loading control. Dot plots are means with 95% confidence interval. *P<0.05 (Mann-Whitney test). **B**, Western blot for Src pTyr^416^ on lysates from the brains of naïve WT mice injected IP with DMSO (n=5 mice) or different doses (0.002 mg/kg (n=6 mice), 0.02 mg/kg (n=4 mice), 2 mg/kg (n=5 mice), 20 mg/kg (n=3 mice)) of AZD 0530. Brains were harvested 6 h after injections. Src was used as the loading control. Dashed line represents the amount of AZD 0530 (0.01465 mg/kg) estimated to be present in the brain after 1 week (10.5 half-lives using the reported half-life of 16 h for AZD 0530 in the brain). Dot plots are means with 95% confidence interval. **C**, representative FACS plots of microglia (gated on CD11b+CD45low cells) in Rhoa^fl/fl^ and Rhoa^fl/fl^:Cx3cr1^CreER+^ mice injected with 10 mg/kg AZD 0530 (one IP per week over 4 weeks) or DMSO. **D**, Rhoa^fl/fl^ and Rhoa^fl/fl^:Cx3cr1^CreER+^ mice injected with 10 mg/kg AZD 0530 (one IP per week over 4 weeks) or DMSO. Representative confocal images of Iba1 immunolabeled cells (asterisks) on hippocampal dentate gyrus are shown. Scale bar: 20 μm. **E,** representative fluorescence images of p65-GFP-transfected (green) pLKO (CT), Rhoa KD or Rhoa KD:Src^Y416F^, Csk KD or Csk KD treated with SKI-1 (500 nM; 24 h) stable N9 microglial cell sub-clones. Nuclei were stained with DAPI (blue). Scale bar: 10 μm. **F**, representative fluorescence images of primary hippocampal neuronal cultures expressing the yellow fluorescent protein variant mVenus were incubated for 24 h with MCM from primary microglia obtained from WT (infected with CT or Rhoa KD lentiviruses) or Tnf KO (infected with Rhoa KD lentiviruses) mice. Scale bar: 50 μm. **G**, representative fluorescence images of primary hippocampal neuronal cultures expressing the yellow fluorescent protein variant mVenus were incubated for 24 h with MCM obtained from pLKO (CT), Rhoa KD, and Rhoa KD + Src^Y416F^ N9 microglial cell sub-clones. In some conditions Rhoa KD microglia were treated with SKI-1 (500 nM; 24 h) or pomalidomide (POM; 500 nM; 24 h). Recombinant Tnf (30 pg/ml; 24 h) was used in CT microglial clones. Scale bar: 50 μm. **H,** Western blotting for synapsin, synaptophysin or PSD-95 on hippocampal tissue lysates from Rhoa^fl/fl^ and Rhoa^fl/fl^:Cx3cr1^CreER+^ mice injected with 10 mg/kg AZD 0530 (one IP per week over 4 weeks) or DMSO (n=8-11 mice). GAPDH was used as the loading control. Dot plots (means and 95% confidence interval) display protein of interest/GAPDH ratio normalized to values obtained from Rhoa^fl/fl^ mice. *P<0.05 (Two-way ANOVA) vs. Rhoa^fl/fl^:Cx3cr1^CreER+^ - DMSO; ^#^P<0.05 (Two-way ANOVA) vs. Rhoa^fl/fl^:Cx3cr1^CreER+^ - AZD. **I and J**, DMSO and AZD 0530 (one IP per week over 4 weeks)-injected Rhoa^fl/fl^:Cx3cr1^CreER+^ mice were evaluated in the NOR test (n=10-11 mice). AZD-injected mice had an overall improvement in recognition index when compared with DMSO-injected Rhoa^fl/fl^:Cx3cr1^CreER+^ mice. Total exploration time was similar in all groups. Dot plots (means and 95% confidence interval) display the NOR parameters after a 4 h delay. *P<0.05 (One-way ANOVA) vs. Rhoa^flfl^ - DMSO; ^#^P<0.05 (One-way ANOVA) vs. Rhoa^flfl^:Cx3cr1^CreER+^ - DMSO. **K**, FACS analysis of microglia in Rhoa^fl/fl^ and Cx3cr1^CreER+^ mice injected with 10 mg/kg AZD 0530 (one IP per week over 4 weeks) or DMSO (n=5-7 mice). Dot plots depict microglia cell counts (mean and 95% confidence interval). Significance was analyzed by Kruskal-Wallis test. **L**, histological confocal analysis of Rhoa^fl/fl^ and Cx3cr1^CreER+^ mice injected with 10 mg/kg AZD 0530 (one IP per week over 4 weeks) or DMSO (n=5-8 mice). vGlut-1 and PSD-95 immunolabeling was analyzed on tissue sections from the hippocampal CA1 region. Dot plots (mean and 95% confidence interval) depicts vGlut-1^+^PSD-95^+^ colocalization puncta normalized to the Rhoa^fl/fl^ - DMSO values. Significance was analyzed by Kruskal-Wallis test. **M and N**, DMSO and AZD 0530 (one IP per week over 4 weeks)-injected Rhoa^fl/fl^ or Cx3cr1^CreER+^ mice were evaluated in the NOR test (n=8-13 mice). No alterations in recognition memory (no significant difference in the discrimination index) was observed between groups. Dot plots (means and 95% confidence interval) display the NOR parameters after a 4 h delay. Significance was analyzed by Kruskal-Wallis test. **O-T**, DMSO and AZD 0530 (one IP per week over 4 weeks)-injected Rhoa^fl/fl^ or Cx3cr1^CreER+^ mice were evaluated in the elevated-plus maze (EPM) and in the open field arena (n=8-9 mice). No alteration in anxiety-related behavior or general locomotor activity (no significant difference in the time spent in the open arms (OA), in the closed arms (CA) or in the center of the maze) was observed between groups. Dot plots are mean and 95% confidence interval and significance was analyzed by Kruskal-Wallis test. **U**, qRT-PCR from the neocortex of Cx3cr1^CreER+^ Rhoa^fl/fl^:Cx3cr1^CreER+^ and Tnf KO mice injected with 50 mg/kg pomalidomide (POM; 3 IPs per week over 4 weeks) or DMSO (n=3-7 mice). Dot plots (means and 95% confidence interval) show the transcript abundance for Tnf normalized to Cx3cr1^CreER+^ – DMSO values. *P<0.05 (Two-way ANOVA) vs. Cx3cr1^CreER+^ - DMSO; ^#^P<0.05 (Two-way ANOVA) vs. Rhoa^flfl^:Cx3cr1^CreER+^ - DMSO; &not significantly different (Two-way ANOVA) vs.Cx3cr1^CreER+^ - POM. **V**, histological confocal analysis in Cx3cr1^CreER+^ and Rhoa^fl/fl^:Cx3cr1^CreER+^ injected with pomalidomide (POM; 3 IPs per week over 4 weeks) or DMSO (n=5 mice per condition). vGlut-1 and PSD-95 immunolabeling was analyzed on tissue sections from the hippocampal CA1 region. Dot plots (mean and 95% confidence interval) depicts vGlut-1^+^PSD-95^+^ colocalization puncta normalized to the Cx3cr1^CreER+^ - DMSO values. *P<0.05 (Two-way ANOVA) vs. Cx3cr1^CreER+^ - DMSO.

**Supplemental Figure 6.**
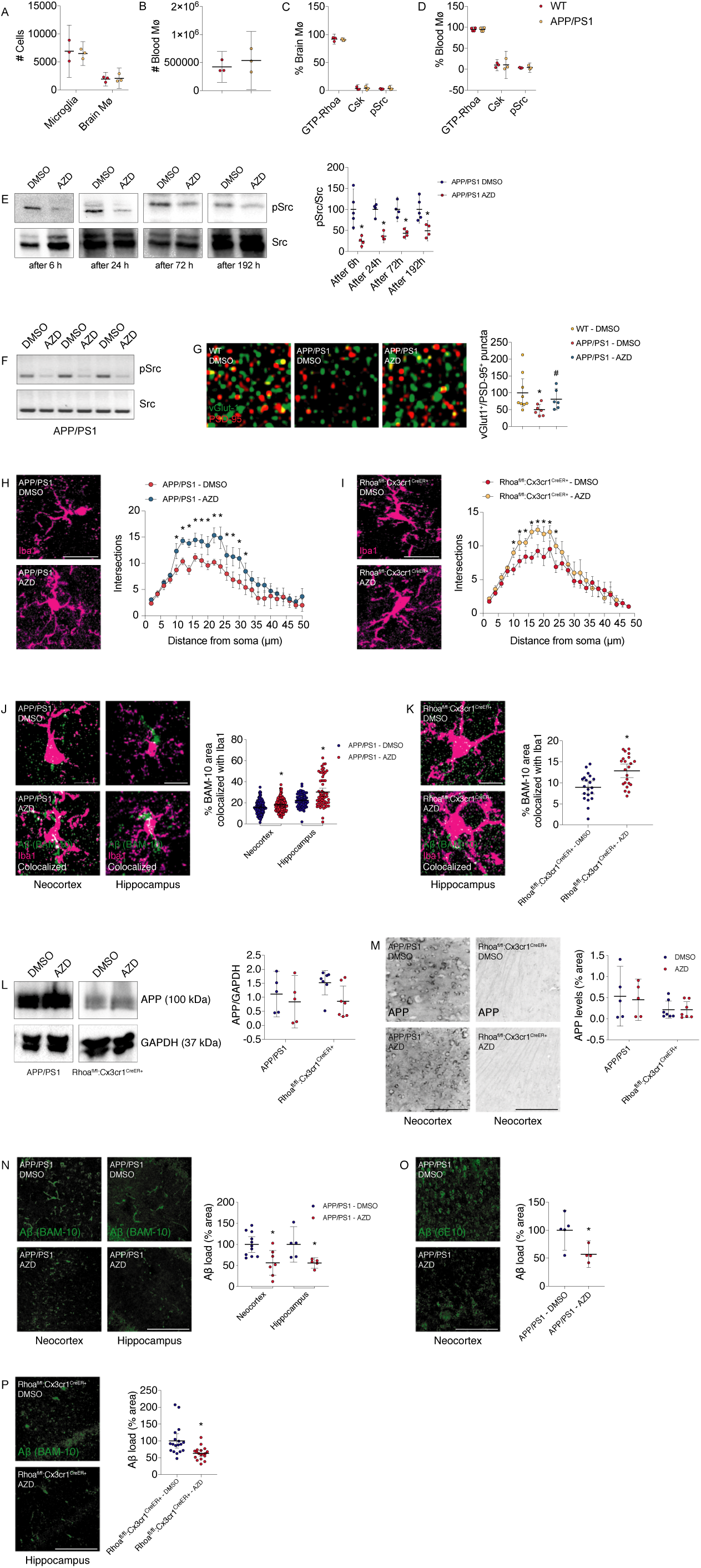
Amyloid affects Rhoa signaling in microglia and inhibition of Src downstream of Rhoa decreases amyloid burden without affecting APP production (Related to Fig. 5 and 6). **A-D**, FACS analysis showing the number of microglia (A), macrophage (A) or blood monocytes (B) in 4-month-old WT or APP/PS1 mice (n=3-4 mice). The percentage of brain macrophages (C) or blood monocytes (D) positive for GTP-Rhoa, Csk or Src pTyr^416^ in 4-month-old WT or APP/PS1 mice are also shown (n=3-4 mice). Dot plots are mean with 95% confidence interval. Significance was analyzed by Mann-Whitney test. **E**, Western blot for Src pTyr^416^ on lysates from the brains of APP/PS1 mice injected IP with a single dose of DMSO (n=4-5 mice per condition) or 10 mg/kg AZD 0530 (n=4 mice per condition). Brains were harvested 6 h, 24 h, 72 h or 196 h after injections. Src was used as the loading control. Dot plots are means with 95% confidence interval. *P<0.05 (Mann-Whitney test). **F**, Western blot for Src pTyr^416^ on lysates from the brains of APP/PS1 mice injected with 20 mg/kg AZD 0530 (one IP per week over 4 weeks) or DMSO (n=3 mice per group). Src was used as the loading control. **G**, histological confocal analysis of APP/PS1 and WT mice injected with 20 mg/kg AZD 0530 (one IP per week over 4 weeks) or DMSO (n=6-7 mice). vGlut-1 (green) and PSD-95 (red) immunolabeling on tissue sections from the hippocampal CA1 region is shown. Dot plots (mean and 95% confidence interval) depicts vGlut-1^+^PSD-95^+^ colocalization puncta normalized to the WT - DMSO values. *P<0.05 (Kruskal-Wallis test) vs. WT - DMSO; #P<0.05 (Kruskal-Wallis test) vs. APP/PS1 - DMSO. Scale bar: 5 μm. **H and I,** histological confocal analysis for Iba1 on tissue sections from the hippocampus of APP/PS1 mice (H) or Rhoa^fl/fl^:Cx3cr1^CreER+^ mice (I) injected with 20 mg/kg or 10 mg/kg AZD 0530 (one IP per week over 4 weeks) or DMSO (n=4 mice for each distance per condition). Dot plots (mean and 95% confidence interval) depicting Sholl analysis for each group is shown. *P<0.05 (Two-way ANOVA comparing DMSO vs. AZD for each distance). Scale bars: 20 μm. **J and K**, histological confocal analysis showing colocalization (white) of Iba1^+^ microglia (magenta) with BAM-10 immunoreactive amyloid deposits (green) on tissue sections from APP/PS1 mice (neocortex and hippocampus; J) or Rhoa^fl/fl^:Cx3cr1^CreER+^ mice (hippocampus; K) injected with 20 mg/kg or 10 mg/kg AZD 0530 (one IP per week over 4 weeks) or DMSO (n= 59-157 cells from 6 mice per condition in J and 20-22 cells from 4 mice per condition in K). Dot plots (mean and 95% confidence interval) show the % area of BAM-10 fluorescence channel colocalized with Iba1. *P<0.05 (One-way ANOVA vs. APP/PS1 - DMSO (J) and Mann-Whitney test (K)). Scale bars: 10 μm. **L**, Western blot for APP on lysates from the brains of APP/PS1 or Rhoa^fl/fl^:Cx3cr1^CreER+^ mice injected with 20 mg/kg or 10 mg/kg AZD 0530 (one IP per week over 4 weeks) or DMSO (n=5 mice per condition in APP/PS1 and 7 mice per condition in Rhoa^fl/fl^:Cx3cr1^CreER+^). GAPDH was used as the loading control. Dot plots are the mean with 95% confidence interval. Significance was analyzed by Two-way ANOVA. **M**, histological confocal analysis showing APP expression levels on neocortical tissue sections from APP/PS1 or Rhoa^fl/fl^:Cx3cr1^CreER+^ mice injected with 20 mg/kg or 10 mg/kg AZD 0530 (one IP per week over 4 weeks) (n=5 mice per condition in APP/PS1 and 7 mice per condition in Rhoa^fl/fl^:Cx3cr1^CreER+^). Dot plots are the mean with 95% confidence interval. Significance was analyzed by Two-way ANOVA. **N and O**, histological confocal analysis of BAM-10 or 6E10 immunoreactive amyloid deposits on tissue sections from APP/PS1 mice (neocortex) injected with 20 mg/kg AZD 0530 (one IP per week over 4 weeks) or DMSO (n=5-11 mice). Dot plots (mean and 95% confidence interval) show the quantification of area occupied by BAM-10 or 6E10 immunoreactive amyloid deposits (Aβ load) normalized to APP/PS1 – DMSO. *P<0.05 (One-way ANOVA vs. APP/PS1 - DMSO (N) and Mann-Whitney test (O)). Scale bars: 200 μm. **P**, histological confocal analysis of BAM-10 immunoreactive amyloid deposits on tissue sections from Rhoa^fl/fl^:Cx3cr1^CreER+^ mice (hippocampus) injected with 10 mg/kg AZD 0530 (one IP per week over 4 weeks) or DMSO (n=18 mice per condition). Dot plots (mean and 95% confidence interval) shows the quantification of area occupied by BAM-10 immunoreactive amyloid deposits (Aβ load) normalized to Rhoa^fl/fl^:Cx3cr1^CreER+^ – DMSO. *P<0.05 (Mann-Whitney test). Scale bar: 200 μm.

